# Unlocking open-access genomic and transcriptomic data: the first bioinformatic exploitation of tunisian durum wheat landraces Chili and Mahmoudi

**DOI:** 10.64898/2026.05.08.723814

**Authors:** Maroua Gdoura-Ben Amor, Nour El Houda Mathlouthi, Imen Belguith, Rania Derouich

## Abstract

Durum wheat (*Triticum turgidum* subsp. *durum*) is a Mediterranean dietary staple threatened by accelerating climate change, yet the genomic basis of adaptation in North African landraces remains poorly characterised. We present the first integrated whole-genome sequencing (WGS) and RNA-seq study of two contrasting Tunisian landraces: humid-adapted Chili and arid-adapted Mahmoudi. From 27,777 high-confidence SNPs, permutation-based F∼ST∼ outlier analysis (1,000 shuffles) identified 46 selection hotspots across six chromosomes, with a peak signal on chromosome 6B (F∼ST∼ = 0.833; *p* = 0.013). Constitutive transcriptome profiling (38,159 expressed genes) revealed 406 expression-divergent observations (|log□FC|□>□1) between landraces. Physical co-localisation analysis confirmed that 99.5% of expression-divergent observations are independent of selection hotspots, implicating trans-regulatory rewiring as the dominant adaptive mechanism. Trans-regulated genes are significantly enriched for disease-resistance (NBS-LRR, RLK, PR; FDR = 1.4 × 10□□) and ubiquitin–proteasome components (FDR = 0.049). Mahmoudi constitutively upregulates ROS-scavenging and dehydrin networks (“store-and-protect”), while Chili elevates aquaporins and transcription factors (“acquire-and-distribute”). These findings identify six chromosomal breeding targets, establish chromosome 6B as a priority fine-mapping locus, and demonstrate that arid-zone adaptation is orchestrated primarily through trans-regulatory stress-network rewiring.

## 1. INTRODUCTION

*Triticum turgidum* subsp. *durum* (2n□=□4x□=□28; AABB genome) is the second most cultivated wheat species worldwide, accounting for approximately 8% of global wheat production and yielding roughly 38 million tons annually (FAOSTAT, 2023).

As the one and only suitable raw material for pasta, couscous, bulgur, and semolina-based staples, durum wheat is the icon of food security and cultural heritage of the Mediterranean basin, North Africa, and the Middle East (Royo et al., 2010). Grown in general under rainfed conditions, present in the most difficult conditions like calcareous soils, erratic rainfall, and high summer temperatures that characterize the Mediterranean climate regime (Royo et al., 2010).

Durum production is being increasingly more vulnerable and threatened by climate change. Shifting precipitation patterns, rising mean temperatures, accelerating drought frequency, and intensifying pathogen pressure collectively erode yield stability in production systems that already operate close to the agronomic margin (Anderson et al., 2020; Arora, 2019; Schlenker and Roberts, 2009). Projection studies indicate that by 2050, key Mediterranean growing regions will experience a 10–25% reduction in water availability during the critical grain-filling period, critically limiting photosynthetic efficiency, nitrogen use, and grain protein deposition (Giorgi and Lionello, 2008; Meddi and Eslamian, 2021).

Sustaining durum production under these conditions demands genetic solutions of a magnitude that the current elite gene pool cannot provide alone.

The most efficient breeding programs have markedly advanced yield potential and disease resistance over the past half-century, but this progress caused a significant reduction in genetic diversity: modern cultivars capture only a fraction of the allelic variation present in their wild and landrace progenitors (Lopes et al., 2015). Unlocking this latent diversity particularly from germplasm adapted to extreme environments is now widely recognized as an essential prerequisite for developing climate-resilient varieties capable of sustaining future production targets (Reynolds and Langridge, 2016).

Over the centuries, farmers saved seeds from their best plants and replanted them every following year and this created populations perfectly tuned to that specific place. Unlike modern pure-line cultivars, landraces are genetically heterogeneous populations that have been continuously shaped by natural selection for local climate, soil, and pathogen pressures, as well as by human selection for agronomic and culinary traits (Zeven, 1998). As a consequence, they harbour substantially higher nucleotide diversity, broader haplotype spectra, and a richer complement of stress-tolerance and resistance alleles than elite germplasm (Cavanagh et al., 2013). These characteristics make landraces invaluable as donors for pre-breeding and molecular introgression programs.

North African durum wheat germplasm constitutes a genetically distinct and underexplored reservoir of adaptive diversity (Robbana et al., 2019). Tunisia, a borderline-situated country between the temperate Mediterranean and the Saharan arid zone, occupies a pivotal position within this reservoir. The country presents one of the steepest agro-ecological gradients in the durum wheat cultivation zone: annual rainfall ranges from 400–700□mm in the humid-to-subhumid north to below 200□mm in the semi-arid to hyper-arid interior, spanning a distance of fewer than 400□km (Latiri et al., 2010; Robbana et al., 2021). Substantial morphological and agronomic diversity among Tunisian landraces have documented over several decades, with populations such as Chili, Mahmoudi, Jenah Zarzoura, and Rommani representing distinct ecotypes adapted to contrasting niches (Ben Krima et al., 2020).

Among this portfolio, Chili and Mahmoudi stand out as the most emblematic and agronomically significant. Both landraces represent opposite ends of the Tunisian environmental gradient. *Chili* originates from the humid-to-subhumid northern regions of Beja, Jendouba, and Siliana (400–700□mm annual rainfall), exhibiting relatively tall stature, broad leaves, and adaptation to heavier clay-loam soils. *Mahmoudi*, by contrast, is one of the oldest documented Tunisian landraces, cultivated since antiquity in the semi-arid to arid interior governorates of Sidi Bouzid, Kairouan, and Gafsa, where annual rainfall falls below 200□mm. Under these conditions, plants face severe and unpredictable water deficit, high solar irradiance, elevated soil temperatures, and intense pathogen pressure during episodic wet seasons. *Mahmoudi* is widely reported to exhibit superior drought tolerance and yield stability under water deficit, and its grain is prized for semolina quality and gluten strength (Chaouachi et al., 2023; Miazzi et al., 2022; Ouaja et al., 2021). Together, these two Tunisian durum wheat landraces represent a natural experiment in local adaptation within a single tetraploid species, shaped by contrasting agro-ecological conditions rather than phylogenetic divergence.

Deeply rooted in Tunisia’s agricultural history, these two landraces are reservoirs of genetic diversity that modern breeding has systematically narrowed, preserving traits linked to superior grain quality, elevated protein content, drought tolerance, and durable disease resistance (Miazzi et al., 2022; Ouaja et al., 2021). Their contrasting adaptive profiles, shaped by opposing ends of the same agro-ecological gradient, make them an ideal natural system for dissecting the genetic and regulatory architecture of environmental adaptation.

Despite their scientific importance, both landraces lacked any comprehensive molecular characterization integrating genome-wide variation with transcriptomic profiling upon recently. This gap was addresses in April 2026 with the launch of the Open Durum Genome Project Tunisia (DurumGPT) consortium (zenodo.org/communities/opendurumgpt/records), which for the first time achieved the complete genome sequencing of both Mahmoudi and Chili using state-of-the-art sequencing technologies (Ayed et al., 2026). By establishing these genomic resources, the consortium positioned this release as a foundation for a new generation of data-driven investigations, encouraging the international research community to explore candidate genes, regulatory networks, and breeding-relevant targets. The present study builds directly upon this initiative. Using these newly available resources, we applied an integrated genomic and transcriptomic approach to investigate the molecular basis of adaptation in these two landraces.

F^ST^-based outlier analysis remains a powerful and interpretable approach to identifying genomic loci under directional selection (Weir and Cockerham, 1984). By detecting allele-frequency divergence that exceeds neutral expectation in a permutation framework, F^ST^ scans can pinpoint discrete chromosomal regions that have been differentially swept in contrasting populations. However, genomic outliers alone cannot resolve the molecular mechanisms through which selection operates. Constitutive transcriptome profiling that is, RNA-seq under ambient, non-stress conditions provides a complementary window onto regulatory divergence that is independent of acute stress responses (Whitehead and Crawford, 2006). The integration of WGS and RNA-seq data within a physical co-localization framework allows a direct test of whether differentially expressed genes reside within, or are genetically uncoupled from, genomic selection hotspots.

This distinction carries fundamental implications for the molecular architecture of adaptation. In the *cis*-regulatory model, adaptive alleles physically linked to their target genes alter local promoter activity, splice-site usage, or mRNA stability, and are therefore co-inherited with altered expression levels (Wittkopp et al., 2004). In the *trans*-regulatory model, distant master regulators transcription factors, chromatin-modifying enzymes, signaling kinases co-ordinate the expression of multiple downstream target genes across the genome without physical linkage to them (Wray, 2007). The distinction matters directly for crop improvement: *cis* variants can be targeted by marker-assisted selection, whereas *trans* loci require eQTL mapping and more complex introgression strategies. Allopolyploid wheat, with its duplicated AABB genome structure, is hypothesized to favor *trans*-regulatory rewiring because the redundancy of homoeologous gene pairs may buffer against deleterious effects of local *cis* mutations, reducing purifying selection at individual regulatory loci (Maccaferri et al., 2019). Whether this mode of regulatory divergence predominates in Tunisian landrace adaptation has not previously been tested.

Against this background, the present study pursues five interconnected objectives. First, we identify genomic loci under directional selection in Chili *versus* Mahmoudi through F^ST^ outlier analysis with permutation-based significance testing on whole-genome SNP data. Second, we characterize constitutive transcriptional divergence between the two landraces across tissues in an exploratory RNA-seq framework. Third, we physically co-localize selection hotspots with expression-divergent observations to distinguish *cis*– from *trans*-regulatory contributions to adaptation. Fourth, we test whether trans-regulatory divergence is functionally coherent and enriched for specific adaptive pathways through gene-set enrichment analysis. Fifth, we identify chromosomal hotspots as prioritized targets for fine-mapping and molecular breeding. Together, these objectives address a critical knowledge gap in the genomics of North African durum wheat and provide the first multi-omics characterization of the molecular basis distinguishing an arid-adapted from a humid-adapted Tunisian landrace.

## 2. MATERIALS AND METHODS

### 2.1 Plant Material and Experimental Design

All data analysed in this study were retrieved from publicly available repositories (see Data Availability). The original plant material comprised two Tunisian durum wheat (*Triticum turgidum L. subsp. durum*) landraces, Chili and Mahmoudi, with contrasting ecoclimatic adaptations. Seeds from a single head per landrace were germinated and grown under standard greenhouse conditions until 10 days post-germination. At that point, ∼5 g of leaf or root tissue was harvested and flash-frozen in liquid nitrogen (Ayed et al., 2026).

Whole-genome sequencing (WGS) was performed on ten accessions (five per landrace; Chili: SRR37204970–SRR37204974; Mahmoudi: SRR37252001–SRR37252005). RNA sequencing (RNA-seq) was performed on four libraries representing root and leaf tissues from each landrace (Chili root: SRR37213950; Chili leaf: SRR37213951; Mahmoudi root: SRR37252182; Mahmoudi leaf: SRR37252142). All sequencing data are deposited under NCBI BioProject PRJNA1420514.

### 2.2 Reference Genome and Annotation

The *T. turgidum* cv. Svevo v1.0 reference genome GCA_900231445.1 (Maccaferri et al., 2019) was used for all analyses. The assembly spans ∼10.5 Gb, with ∼9.4 Gb anchored to 14 pseudochromosomes (LT934111.1–LT934124.1; chromosomes 1A–7B). Gene models were retrieved from NCBI in EMBL GFF3 format, comprising 63,863 features, of which 62,814 carried Automated Human Readable Description (AHRD) product annotations.

### 2.3 WGS Pre-processing, Alignment, and Variant Calling

Raw reads were assessed for quality with FastQC and trimmed with Trimmomatic v0.39. Short reads were aligned to the reference genome using minimap2 v2.30 (––sr) via the Galaxy Europe platform (usegalaxy.eu). Duplicate reads were marked with Picard MarkDuplicates v3.1.1.0 (optical duplicate pixel distance = 2,500). Variant calling was performed with FreeBayes v1.3.10 under tetraploid assumptions (ploidy = 4), with the following parameters: ––min-mapping-quality 30, ––min-base-quality 20, ––min-coverage 10, ––min-alternate-fraction 0.1, ––min-alternate-count 2, and ––theta 0.001.

### 2.4 Variant Filtering, Annotation, and Merging

Variant quality filtering required: QUAL ≥ 20, DP ≥ 5, MQM ≥ 30, SAF > 0, and SAR > 0 (bidirectional strand representation), and TYPE = snp. Functional annotation was conducted with SnpEff v5.4c using a custom Svevo v1.0 database; only variants with HIGH or MODERATE impact were retained. Merging across samples was performed with bcftools merge v1.19 (––missing-to-ref), yielding 27,777 high-confidence SNP sites. For population genomic analyses, tetraploid genotypes were converted to diploid equivalents: 0 alternate alleles → 0/0 (homozygous reference); 1-3 alternate alleles → 0/1 (heterozygous); 4 alternate alleles → 1/1 (homozygous alternate).

### 2.5 Population Genomics, FST Permutation, and PCA

Genome-wide differentiation was estimated using the Weir–Cockerham FST estimator implemented in VCFtools v0.1.16. For visualisation, FST was calculated in 100-kb windows with a 10-kb step size; for outlier detection, 10-kb windows with a 1-kb step size were used, retaining only windows with ≥ 3 variants. Empirical significance was assessed via permutation testing: 1,000 label-shuffle permutations were performed by randomly reassigning the ten samples into two groups of five, recalculating genome-wide FST, and recording the maximum FST per permutation. The empirical P-value was calculated as (number of permutations with maximum FST ≥ observed + 1) / (Nperm + 1), with significance defined at P < 0.05. The distribution of permuted maximum FST values is shown in Figure S2. Nucleotide diversity (π) and Tajima’s D were calculated in 100-kb windows per population. Principal component analysis (PCA) was conducted in PLINK v2.00a6 on biallelic SNPs with minor allele frequency (MAF) ≥ 15% (n = 9 after excluding SRR37204972; SRR37204972 was excluded from PCA due to elevated variant count (16,603), consistent with an alternative processing path, suggesting technical heterogeneity; the sample was retained for FST and diversity analyses). No linkage disequilibrium (LD) pruning was performed because PLINK v2 requires n ≥ 50 for ––indep-pairwise; PCA was therefore conducted on the biallelic, MAF ≥ 15% SNP set after excluding SRR37204972.

### 2.6 Physical Co-localisation

Physical co-localisation between FST outlier windows and gene bodies was determined using an in-house Python implementation of interval intersection (analogous to bedtools intersect –wa – wb). FST outlier windows were extended ±50 kb from their boundaries and intersected with the 63,863 Svevo v1.0 GFF3 gene features. Overlapping gene IDs were cross-referenced against 406 expression observations.

### 2.7 RNA-seq Pre-processing, Alignment, and Quantification

Raw reads were assessed for quality with FastQC and trimmed with Trimmomatic v0.39. Per-library alignment statistics are provided in Table S4. Reads were aligned with HISAT2 v2.2.1. Alignments were sorted and indexed with SAMtools v1.21. Gene-level quantification was performed with featureCounts (Subread v2.0.6) using parameters –t gene –g ID –F GFF –p ––countReadPairs, with multi-mapping reads discarded to ensure a conservative approach to homeolog ambiguity in allotetraploid wheat.

### 2.8 Constitutive Transcriptome Profiling

Differential expression analysis was performed with DESeq2 v1.44 in R v4.6.0 using an additive model (design = ∼ landrace + tissue). A critical caveat applies: the experimental design comprises n = 1 per landrace × tissue combination, providing zero within-group biological replication for the landrace contrast. Consequently, gene-wise dispersion relies entirely on the global mean–dispersion trend, and no gene-specific variance estimate is available. We therefore frame this analysis as exploratory observation ranking rather than formal hypothesis testing; adjusted P-values are not reported as significance thresholds for the landrace contrast, and Type I error control is not achievable without replication.

Genes with fewer than 10 counts in fewer than 2 samples were excluded, retaining 38,159 genes. Of these, 2,220 genes were subsequently excluded by DESeq2 internal filtering (zero-count in at least one sample across the n□=□1 design), leaving 35,939 genes for differential expression testing. Effect sizes were estimated using adaptive shrinkage (lfcShrink type = “ashr”) (Stephens, 2017). A biological relevance threshold of |ashr-shrunk log□fold change| > 1 was applied. To mitigate high-variance, low-reliability observations inherent to the n = 1 design, a secondary noise filter requiring DESeq2-adjusted P < 0.05 (from the global mean–dispersion model) was additionally imposed. All 406 retained observations satisfied both criteria (|ashr-shrunk log□FC| > 1 and P < 0.05), though we emphasise that P values serve solely as a noise-filtering criterion. The initial set comprised 547 observations; after ashr shrinkage, 406 directional observations were retained (204 Chili-elevated; 202 Mahmoudi-elevated).

Because the additive model constrains the landrace effect to be identical across tissues, cross-tissue consistency is 100% by mathematical construction, a property that does not constitute biological replication. Genuine tissue-specific effects would require n ≥ 3 per group per tissue. We therefore term these 406 observations expression-divergent observations rather than differentially expressed genes; they require biological replication (n ≥ 3) and qRT–PCR validation. Cross-tissue consistency verification is provided in Table S7. The variance-stabilising transformation (VST; meaning dispersion is estimated without conditioning on the design matrix, ensuring an unbiased visual representation) was applied only to PCA and heatmap visualisation, not to differential testing.

### 2.9 Functional Classification

AHRD product annotations were manually curated into 16 biological categories (6 of 16 categories are shown in Table 1; full enrichment statistics for all 12 categories are in Table S8). Of these, 12 categories containing sufficient observations were tested for enrichment relative to the expressed transcriptome background using Fisher’s exact test (one-tailed; BH FDR correction); four minor categories were excluded due to insufficient observations for reliable enrichment testing.

**Table 1:** Major functional categories among 406 expression-divergent candidates.

| Functional category | Total (n) | Chili-up | Mahmoudi-up | Mean $\vert \log_2 \text{FC}\vert $ | Cross-ref (n) |
| --- | --- | --- | --- | --- | --- |
| Disease resistance (NBS-LRR, | 91 | 36 | 55 | ~4.1 | 91 |

| RLK, PR) |  |  |  |  |  |
| --- | --- | --- | --- | --- | --- |
| Drought / ABA signalling | 38 | 17 | 21 | ~2.3 | 38 |
| ROS scavenging | 32 | 1 | 31 | ~2.8 | 32 |
| Grain quality (glutenins,<br>gliadins) | 18 | 10 | 8 | ~2.1 | 18 |
| Heat stress / chaperones | 12 | 8 | 4 | ~1.9 | 0 |
| Transcription factors | 28 | 25 | 3 | ~1.8 | 0 |
| TOTAL (above categories) | 219 | 97 | 122 | - | 179 |

### 2.10 Functional Enrichment of Trans-regulatory Observations

To test whether 404 trans-regulatory observations (406 total minus 2 co-localised genes not within ±50 kb of FST outlier windows) represent functionally coherent stress adaptation or random transcriptional noise, we performed Fisher’s exact test (one-tailed alternative = “greater”) against a background of 35,535 non-divergent expressed genes (retained from 35,939 DESeq2-tested genes after excluding the 404 trans-regulatory observations). AHRD product descriptions were used for annotation, with 12 enrichment-tested biological categories (of 16 curated; Table S8). Multiple testing correction was performed using the Benjamini–Hochberg false discovery rate (FDR), with significance defined at FDR < 0.05.

Sensitivity analyses included: (a) background = all 35,939 expressed genes versus 35,535 non-divergent genes; (b) trans set = 404 genes (with 2 cis genes excluded from the 406 total); and (c) 10-category collapsed classification versus 12 categories. No formal GO or KEGG annotation is available for Svevo v1.0; the AHRD keyword-based approach is therefore approximate. Sensitivity analyses are documented in enrichment_sensitivity.txt; distance threshold and LD analyses are detailed in Supplementary Notes S1 and S2, respectively.

### 2.11 Structural prediction and cis-regulatory variant screening

Landrace-fixed protein-coding variants were identified from SnpEff-annotated VCFs (v5.4c, Svevo v1.0 database) using a strict fixation filter: alternate allele count ≥8 of 20 tetraploid alleles (≥4 of 5 accessions) in one landrace and ≤2 of 20 in the other. All three landrace-fixed variants reported in §3.12 have 0 alternate alleles in the opposite landrace; the operational ≤2 threshold captures near-complete fixation without discarding genuine near-fixation events. Stop-gained, frameshift, and splice-site variants were captured using an expanded HGVS.p parser (not limited to missense patterns) to avoid silent dropping of truncating variants. The ±5 kb cis-regulatory window (gene body ±5 kb upstream and downstream) was scanned in the unfiltered merged diploid VCF (durum_merged_diploid.vcf.gz; 27,777 high-confidence SNPs, post-FreeBayes quality filtering with QUAL ≥ 20, DP ≥ 5, MQM ≥ 30) for landrace-biased MODIFIER and LOW-impact variants (upstream, UTR, intronic, synonymous). This window does not capture distal enhancers or chromatin-level regulatory elements. AlphaFold 3 Server (alphafoldserver.com) was used to predict the monomeric structure of the Svevo reference allele of TRITD_5Av1G148980; the prediction cannot capture NLR oligomerisation states or effector-bound conformations. Per-residue pLDDT and predicted TM-score (pTM) were extracted from the .cif output. Nonsense-mediated decay was not formally predicted; the NMD status of the truncated allele was inferred from RNA-seq read-count abundance. qRT-PCR with allele-specific primers and ribosome profiling will be required to formally separate transcriptional regulation from NMD efficiency. Structural visualisation was performed in PyMOL v2.5 (Schrödinger) with the Svevo prediction coloured by domain and the Mahmoudi truncation boundary annotated at residue 384. Per-accession genotypes in Figure 5e are reported on the original tetraploid scale (0/0/0/0 or 1/1/1/1) before diploid conversion for population analyses. The merged VCF was generated with bcftools merge ––missing-to-ref, which converts missing genotypes to homozygous reference; this may introduce false-negative calls at low-coverage loci.

## 3. RESULTS

### 3.1 WGS Data Quality and Variant Landscape

After quality filtering, the final variant set comprised 27,777 high-confidence SNP sites. Per-sample variant counts ranged from 1,532 to 4,313 (median = 2,232; Table S9). One Chili accession (SRR37204972) exhibited 16,603 variants, consistent with an alternative processing path; this sample was retained for FST and diversity analyses but excluded from PCA

### 3.2 Population Structure Resolves Landraces

Principal component analysis of 3,654 biallelic SNPs (MAF ≥ 15%) revealed partial landrace separation (Figure 1). PC1 explained 25.4% of the variance and showed partial discrimination between landraces, while PC2 explained 15.0% and provided additional resolution. Chili accessions clustered predominantly in the positive PC1 and PC2 space (−0.08 to +0.25), whereas Mahmoudi accessions occupied the negative PC2 space (−0.276 to −0.456). One Mahmoudi accession (SRR37252002) was a clear outlier on PC1 (−0.907), possibly reflecting elevated heterozygosity or a structural variant. The partial separation is consistent with landrace identity but indicates substantial shared ancestral variation.

**Figure 1:**
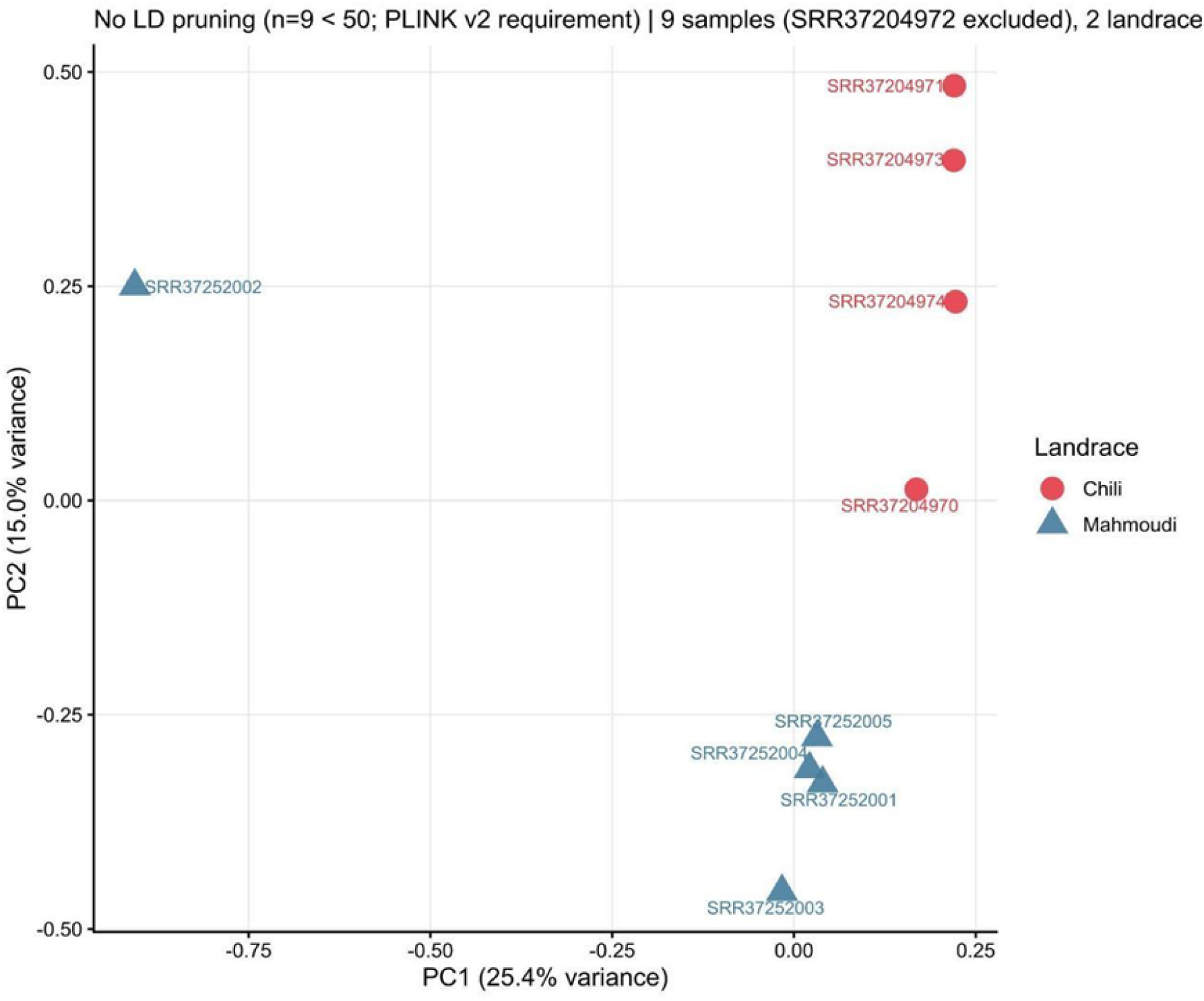
Population structure resolves Tunisian durum wheat landraces from whole-genome sequencing data. Principal component analysis of 3,654 biallelic SNPs (MAF ≥ 15%) from n = 9 accessions, excluding SRR37204972. No linkage disequilibrium pruning was performed (PLINK v2 requires n ≥ 50). PC1 explains 25.4% of the variance and shows partial landrace separation; PC2 explains 15.0% and provides additional resolution. Chili accessions (red circles) cluster in positive PC1 and PC2 space. Mahmoudi accessions (blue triangles) occupy negative PC2 space. SRR37252002 is an outlier on PC1 (−0.907), possibly reflecting elevated heterozygosity or a structural variant. Partial separation is consistent with landrace identity but indicates substantial shared ancestral variation.

### 3.3 Genome-Wide FST Reveals Selection Hotspots

Genome-wide differentiation analysis identified 46 empirically significant outlier windows with FST ≥ 0.50 (Figure 2). The strongest signal occurred on chromosome 6B ( FST = 0.833, empirical P = 0.013), indicating near-complete allele fixation. Across 7,203 tested 100-kb windows (≥3 variants), mean weighted FST was 0.083, and median FST was 0.050, indicating modest but genuine genomic differentiation.

**Figure 2:**
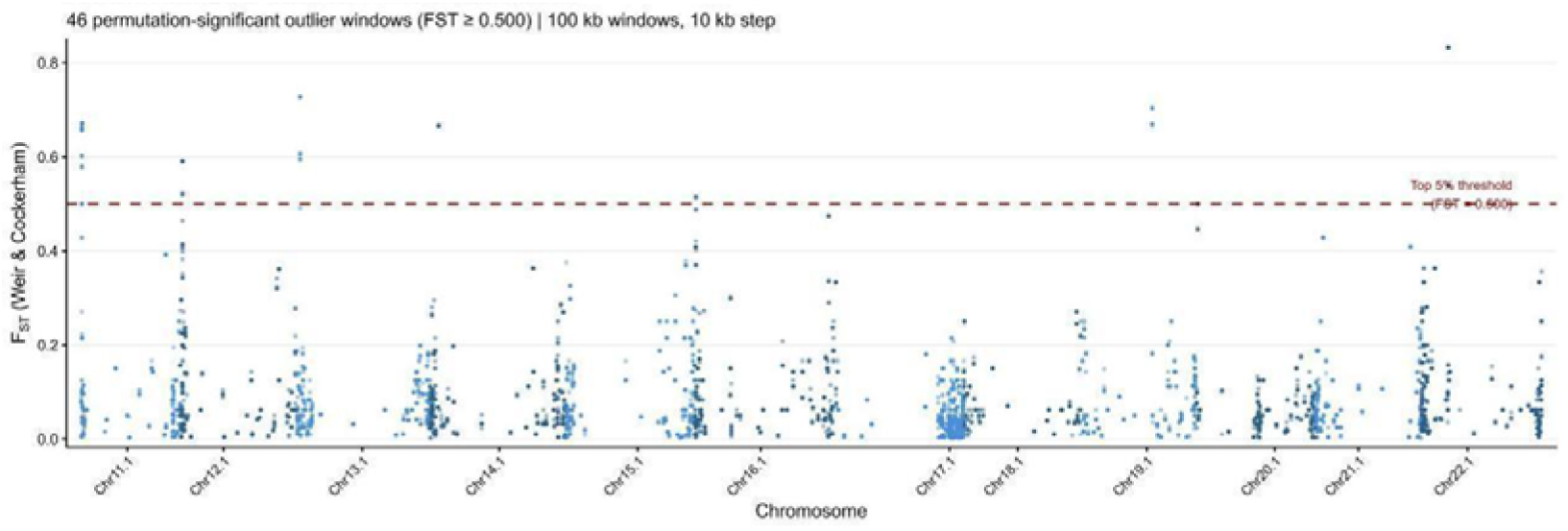
Genome-wide differentiation reveals six chromosomal selection hotspots. Weighted FST (Weir–Cockerham) in 100-kb windows with 10-kb step across all 14 pseudochromosomes of Triticum turgidum cv. Svevo v1.0. 46 permutation-significant outlier windows (FST ≥ 0.50, empirical P < 0.05 from 1,000 label-shuffle permutations) are distributed across chromosome 1A (n = 9), 1B (n = 1), 2A (n = 10), 2B (n = 6), 5A (n = 10), and 6B (n = 10). The horizontal dashed line marks FST = 0.50 (95th percentile of permuted maximum FST). The maximum signal occurs on chromosome 6B (FST = 0.833), which equalled the 99th percentile of the permuted null distribution (empirical P = 0.013), indicating near-complete allele fixation under strong directional selection. X-axis labels use NCBI accession-derived identifiers from the Svevo v1.0 reference assembly; correspondence to standard wheat chromosome nomenclature is: Chr11.1 = 1A, Chr12.1 = 1B, Chr13.1 = 2A, Chr14.1 = 2B, Chr15.1 = 3A, Chr16.1 = 3B, Chr17.1 = 4A, Chr18.1 = 4B, Chr19.1 = 5A, Chr20.1 = 5B, Chr21.1 = 6A, Chr22.1 = 6B, Chr23.1 = 7A, Chr24.1 = 7B.

The 46 outlier windows (10-kb analysis) were distributed across six chromosomes: 1A (n = 9), 1B (n = 1), 2A (n = 10), 2B (n = 6), 5A (n = 10), and 6B (n = 10). The 6B hotspot at ∼154.4 Mb (LT934122.1) spanned ∼30 kb and exceeded the 99th percentile of neutral expectations (permuted maximum FST distribution; Figure S2). Permutation testing (1,000 label-shuffle permutations) confirmed that all 46 outliers were significant at P < 0.05; the 95th and 99th percentiles of the permuted maximum FST distribution were 0.500 and 0.833, respectively. Per-chromosome summary statistics are provided in Table S3.

### 3.4 Nucleotide Diversity and Shared Demographic History

Nucleotide diversity (π) and Tajima’s D distributions were comparable between landraces (Figure 3; Table S3). Genome-wide median π in 100-kb windows (≥3 variants) was 1.00 × 10□□ for Chili (n = 13,794 windows) and 1.11 × 10□□ for Mahmoudi (n = 12,682 windows), indicating broadly similar diversity levels. Chili π exceeded Mahmoudi π only on chromosomes 5B, 6B, and 7B. Tajima’s D was uniformly negative for both landraces (median = −1.11; Chili: 99,587 windows; Mahmoudi: 99,520 windows), consistent with a shared demographic history of expansion or bottleneck. This supports the interpretation that differentiation is driven by selection at discrete loci rather than genome-wide drift.

**Figure 3:**
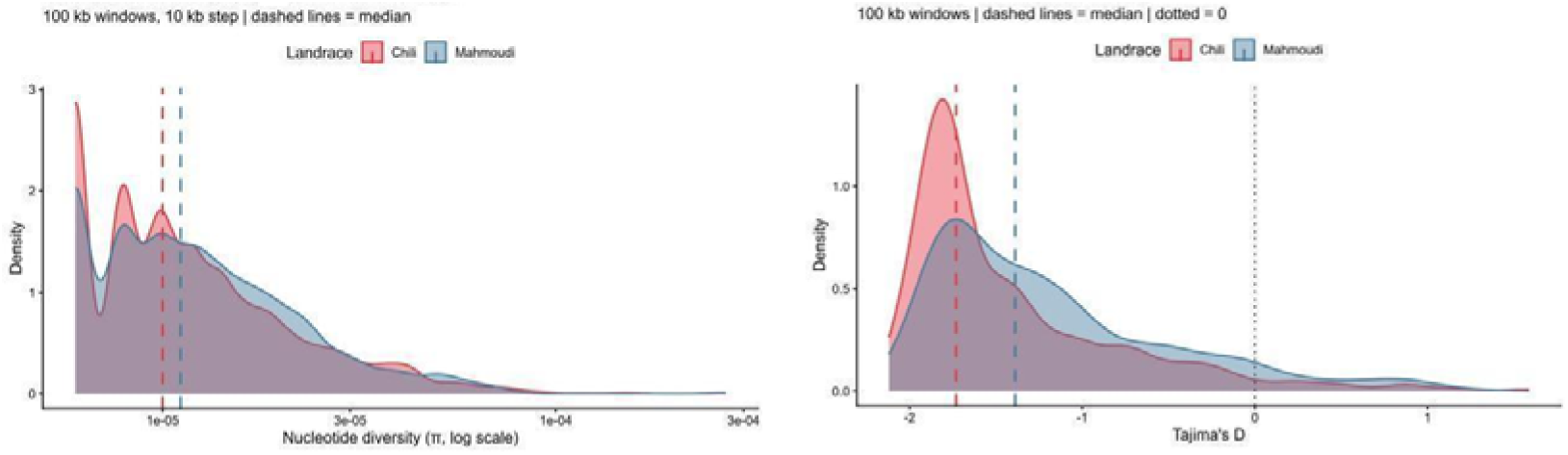
Comparable nucleotide diversity and shared demographic history between landraces. (a) Kernel density estimates of nucleotide diversity (π) in 100-kb windows (≥ 3 variants). Genome-wide median π was 1.00 × 10LL for Chili (n = 13,794 windows) and 1.11 × 10LL for Mahmoudi (n = 12,682 windows), indicating broadly similar diversity levels. Note: The x-axis is on a log scale. (b) Tajima’s D distributions with median = −1.11 for both landraces (Chili: 99,587 windows; Mahmoudi: 99,520 windows). The uniformly negative distribution indicates a shared demographic history of expansion or bottleneck, supporting the interpretation that differentiation is driven by selection at discrete loci rather than genome-wide drift.

### 3.5 Physical Co-Localisation of Selection and Expression Signals

Physical co-localisation analysis intersected FST outlier windows (±50 kb) with 406 expression-divergent observations. Only 2 genes (0.5%) were physically co-localised (Figure 4; Table S6): TRITD_5Av1G148950 (chloroplast movement gene, Chili-elevated) and TRITD_5Av1G148980 (TIR-NBS-LRR disease resistance gene, Mahmoudi-elevated), both on chromosome 5A. The remaining 404 observations (99.5%) were beyond ±50 kb, indicating that the vast majority of expression divergence is not physically linked to selection hotspots.

**Figure 4:**
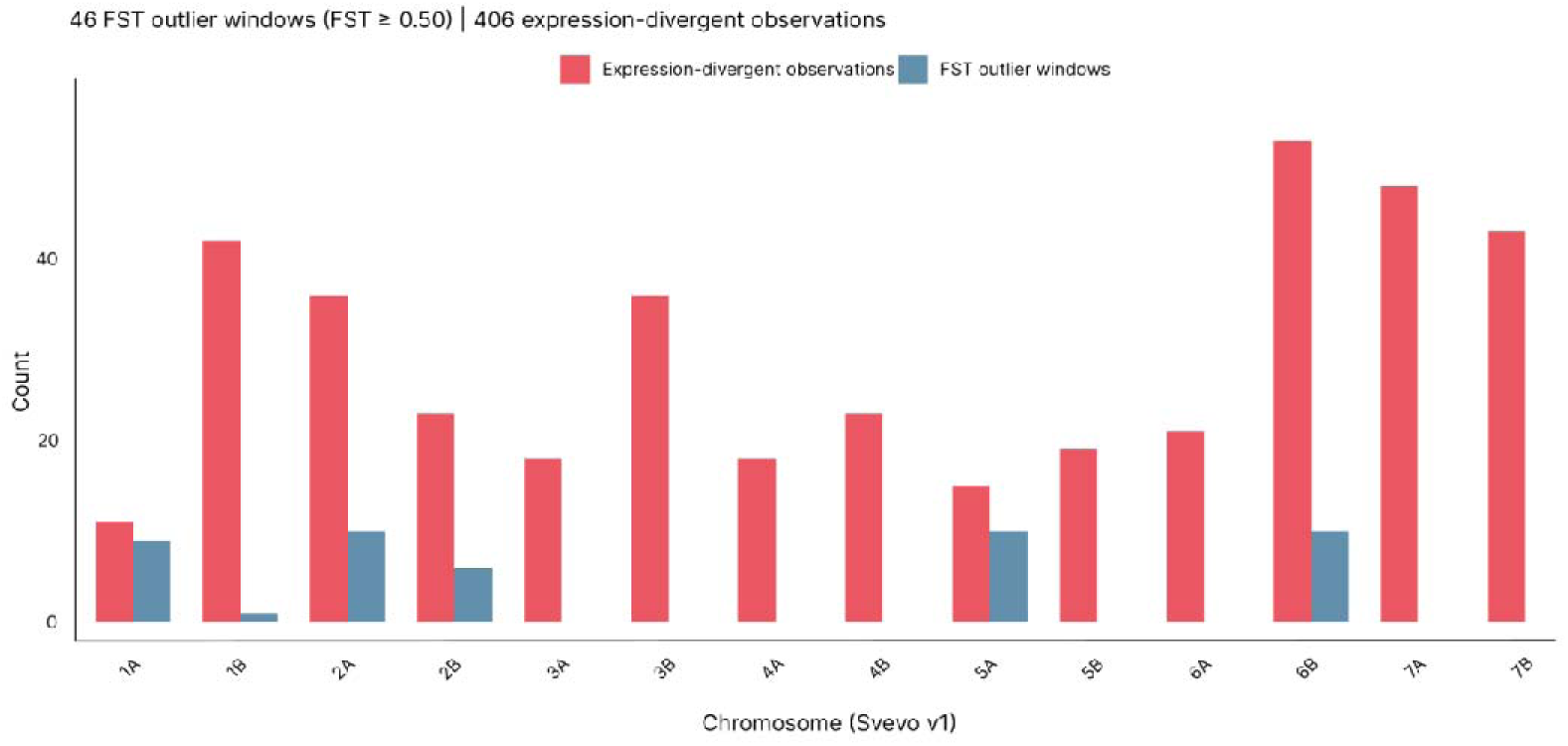
Physical co-localisation of selection hotspots and expression divergence. Bar chart showing the number of expression-divergent observations per chromosome (n = 406 total). Asterisks denote chromosomes harbouring FST outlier windows ( FST ≥ 0.50). Chromosome 6B combines the strongest selection signal ( FST = 0.833) with the largest expression cluster (53 observations), yet zero of these 53 observations physically overlap the hotspot at ±50 kb, ±500 kb, or ±1 Mb. Only 2 genes (0.5%) physically co-localise with FST outliers at ±50 kb: TRITD_5Av1G148950 (chloroplast movement, Chili-elevated) and TRITD_5Av1G148980 (TIR-NBS-LRR disease resistance, Mahmoudi-elevated), both on chromosome 5A. The predominance of chromosomal co-occurrence without physical overlap supports trans-regulatory divergence as the primary mechanism of constitutive expression differentiation. X-axis uses standard wheat chromosome nomenclature (1A–7B); correspondence to Svevo v1.0 NCBI accession-derived identifiers is: Chr11.1 = 1A, Chr12.1 = 1B, Chr13.1 = 2A, Chr14.1 = 2B, Chr15.1 = 3A, Chr16.1 = 3B, Chr17.1 = 4A, Chr18.1 = 4B, Chr19.1 = 5A, Chr20.1 = 5B, Chr21.1 = 6A, Chr22.1 = 6B, Chr23.1 = 7A, Chr24.1 = 7B.

**Figure 5:**
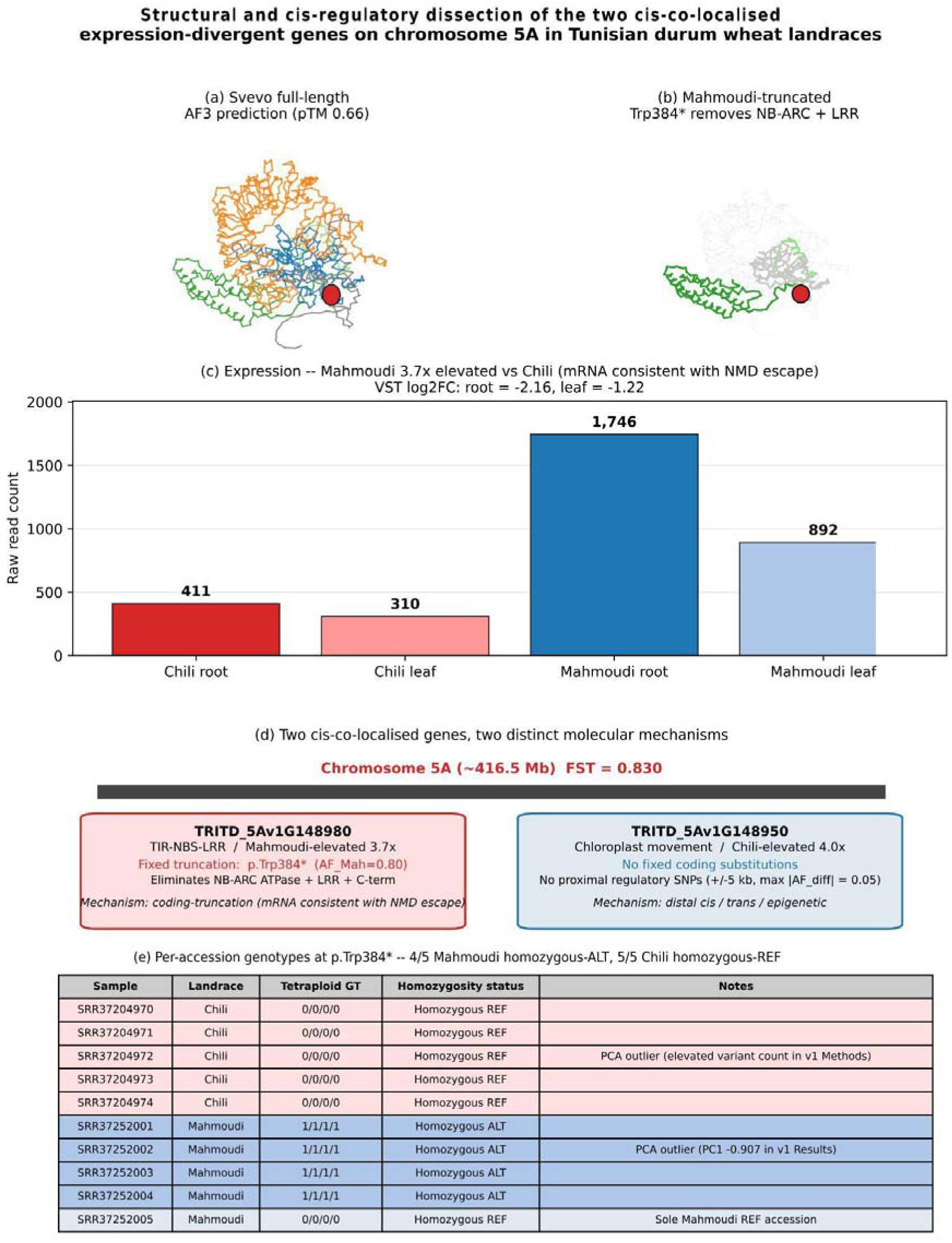
Structural and cis-regulatory dissection of the two cis-co-localised expression-divergent genes on chromosome 5A in Tunisian durum wheat landraces. (a) Svevo full-length TIR-NBS-LRR (TRITD_5Av1G148980) AlphaFold 3 prediction (pTM 0.66), coloured by domain: TIR (green, 1–160), linker (light green, 160–200), NB-ARC (blue, 200–500), LRR (orange, 500–1,100), C-terminal (grey, 1,100–1,224). Residue 384 (Trp) marked with red sphere indicating truncation site. (b) Mahmoudi-truncated structure: residues 384––1,224 masked in transparent grey, showing retained N-terminal fragment (TIR + N-terminal NB-ARC, residues 1––383). (c) Expression bar chart using VST-normalised counts for TRITD_5Av1G148980 (raw counts in parentheses): Chili root (411), Chili leaf (310), Mahmoudi root (1,746), Mahmoudi leaf (892), showing 3.7× Mahmoudi elevation (mRNA consistent with NMD escape). VST log□FC: root = −2.16, leaf = −1.22. (d) Schematic of chromosome 5A hotspot (∼416.5 Mb, FST = 0.830) showing two cis-co-localised genes with divergent mechanisms: TRITD_5Av1G148980 (coding truncation, Mahmoudi-elevated) and TRITD_5Av1G148950 (no proximal sequence divergence, Chili-elevated). (e) Per-accession genotype table for p.Trp384* (10 accessions: 5 Chili homozygous REF, 4 Mahmoudi homozygous ALT, 1 Mahmoudi homozygous REF).

Chromosome 6B exemplified this spatial decoupling: although 53 expression-divergent observations mapped to 6B, the largest chromosomal cluster, none were within ±50 kb of the 6B FST hotspot. Similarly, chromosomes 7A (n = 48 observations) and 7B (n = 43) harboured expression divergence but no FST outliers, implying entirely trans-regulatory mechanisms. Sensitivity analyses tested distance threshold robustness. At ±500 kb, 6 observations (1.5%) were co-localised, including 4 genes beyond the ±50 kb set; at ±1 Mb, the count plateaued at 6 (1.5%), confirming that all physically linked genes were captured within 500 kb. Thus, trans-regulatory divergence (≥98.5%) predominates regardless of distance threshold.

LD analysis on chromosome 6B supported genetic independence of the expression cluster from the selection signal. Of 50 SNPs within ±5 Mb of the lead SNP (sparse coverage due to the small WGS panel), zero of the 53 chromosome-6B observations showed r² > 0.2 with the lead SNP, and the median r² was ≤0.25 across distance bins. Rapid LD decay confirms that the 53 observations are genetically independent of the 6B hotspot; physical separation (>1 Mb) remains the primary evidence for trans-regulatory divergence (Supplementary Note S2).

### 3.6 RNA-Seq Data Quality

RNA-seq alignment and quantification metrics confirmed high data quality (Table S4). The four libraries yielded 132,711,276 total read pairs (range: 30.99–36.40 million per library). Alignment rates were 93.08–94.70%, concordant unique alignment was 83.0–84.6%, and featureCounts assignment was 70.5–75.0%. Multi-mapping reads (20.2–24.3%) were discarded to mitigate homeolog ambiguity. After filtering, 38,159 genes were retained. The mean-dispersion trend was well fitted (Figure S1).

### 3.7 Transcriptome Structure

PCA of variance-stabilised counts (top 500 genes by variance) confirmed biological coherence and absence of technical confounding (Figure S5a). PC1 explained 99.5% of the variance and corresponded to tissue identity (root vs. leaf), while PC2 explained 0.3% and corresponded to landrace identity. Tissue dominates transcriptional variance; the landrace signal is detectable but minor. Four-way sample separation demonstrates that the landrace contrast is embedded within a robust tissue-driven structure.

### 3.8 Constitutive Transcriptome Divergence

Complete tissue differential expression results are provided in Table S2. Constitutive transcriptome divergence between landraces is visualised in Figure S5b (MA plot), Figure S5d (heatmap), Figure S5e (functional categories), and Table 1 (category statistics). 406 expression-divergent observations were identified (|ashr-shrunk log□ fold change| > 1): 204 Chili-elevated and 202 Mahmoudi-elevated. The absolute log□ fold change range was 1.0–8.3 (median ≈ 1.7). The initial set comprised 547 observations, reduced to 406 after adaptive shrinkage. All 406 showed 100% cross-tissue consistency, a mathematical property of the additive model constraint (n = 1 per landrace × tissue; see Materials and Methods).

Functional classification of the 406 observations (Table S1) revealed major adaptive categories. Disease resistance comprised 91 observations (36 Chili-elevated, 55 Mahmoudi-elevated; mean |log□FC| ≈ 4.1). Drought and ABA signalling comprised 38 (17 Chili, 21 Mahmoudi; mean |log□FC| ≈ 2.3). ROS scavenging comprised 32 (1 Chili, 31 Mahmoudi; 96.9% Mahmoudi bias; mean |log□FC| ≈ 2.8). Grain quality proteins (glutenins, gliadins) comprised 18 (10 Chili, 8 Mahmoudi; mean |log□FC| ≈ 2.1). Heat stress and chaperones comprised 12 (8 Chili, 4 Mahmoudi; mean |log□FC| ≈ 1.9). Transcription factors comprised 28 (25 Chili, 3 Mahmoudi; mean |log□FC| ≈ 1.8). These six categories accounted for 219 of 406 observations; the remaining 187 fell into minor categories. Note: Manual-curation counts in Table 1 (91 disease resistance, 32 ROS scavenging, 38 drought / ABA signalling) include genes with partial keyword matches; AHRD strict-keyword counts (67, 6, 4) are used for the statistical enrichment in Table S8. These seedling storage reserves are not indicative of mature grain phenotype.

### 3.9 Cross-Study Validation

Cross-study validation against nine published wheat stress studies (Figure S5f; Table S5) showed that 160 of 406 observations (39.4%) matched gene families reported in these studies. The deduplicated category breakdown was: drought/dehydrin/LEA genes (n = 38), ROS/antioxidant genes (n = 32), stem rust/NBS-LRR genes (n = 91), abiotic stress genes (n = 12), and root architecture genes (n = 8; raw category sum = 181; deduplicated total = 160, with 21 observations assigned to multiple categories). The remaining 246 observations (60.6%) showed no published wheat stress counterpart, representing potentially novel Tunisian-landrace-specific regulatory loci.

### 3.10 Functional Enrichment Confirms Trans-Regulatory Observations Target Stress Pathways

To test whether the 404 trans-regulatory observations represent functional coherence or random noise, we performed Fisher’s exact test (12 AHRD-keyword categories; background = 35,535 non-divergent expressed genes; Benjamini–Hochberg FDR correction). Disease resistance (including NBS-LRR, RLK, and PR proteins) was strongly enriched: 67 of 404 trans observations (16.6%) vs. 2,512 of 35,535 background genes (7.1%); odds ratio = 2.6; FDR = 1.4 × 10□□. Of the 91 total disease resistance observations, 90 are trans-regulatory (>1 Mb from any FST hotspot); the single cis-co-localised example is TRITD_5Av1G148980 (TIR-NBS-LRR, chromosome 5A, Mahmoudi-elevated). Note: the enrichment analysis identified 67 disease resistance genes by AHRD keyword matching, a subset of the manually curated 91. Ubiquitin-proteasome system genes were also significantly enriched: 36 of 404 (8.9%) vs. 2,064 of 35,535 (5.8%); odds ratio = 1.6; FDR = 0.049. No significant depletions were observed (Figure S3; Table S8).

Sensitivity analyses confirmed robustness. Analysis (a) (background = all 35,939 expressed genes): disease resistance remained significant (odds ratio = 2.6; FDR = 2.6 × 10□□); ubiquitin-proteasome became marginally non-significant (FDR = 0.054). Analysis (b) (trans set = 404 with 2 cis genes excluded): disease resistance was identical to the original (FDR = 1.4 × 10□□); ubiquitin-proteasome was identical (FDR = 0.049). Analysis (c) (10-category collapsed classification): disease resistance strengthened (odds ratio = 2.7; FDR = 5.8 × 10□¹¹); ubiquitin-proteasome remained robust (FDR = 0.037).

Disease resistance enrichment is highly robust across all scenarios. Ubiquitin-proteasome enrichment is borderline (FDR 0.037–0.054) and should be reported with the caveat that significance is scenario-dependent.

These results demonstrate that trans-regulatory divergence is non-random and targets the same adaptive pathways, disease resistance and stress proteostasis, identified by FST selection analysis. The multi-omic convergence of genomic selection and trans-regulation on common functional modules suggests that selection may act on master trans-regulators that coordinate expression of dispersed adaptive gene batteries.

### 3.11 Interactive visualisation of population-genomic and transcriptomic divergence

To make the dataset more accessible to the broader community, we developed TuniTritica, a browser-based interactive tool. It was built as a single self-contained HTML file using Plotly.js, with no server-side dependencies or installation required. The interface exposes all 406 expression-divergent observations and 46 FST outlier windows through dynamic tables that link to chromosome-wide population-genomic charts. Users filter by chromosome, landrace bias, |log□FC|, or AHRD functional category, then click any chromosome to retrieve windowed FST, nucleotide diversity (π), Tajima’s D, and empirical permutation P-values. All underlying data tables are exportable as CSV. The webapp is deployed at https://nour0810.github.io/TuniTritica/; the complete codebase, processed data, and analysis scripts are version-controlled at https://github.com/nour0810/TuniTritica.

### 3.12 Structural and cis-regulatory dissection of the two cis-co-localised expression-divergent candidates

To distinguish coding from regulatory drivers of the two cis-co-localised expression-divergent genes on chromosome 5A, we screened all 406 expression-divergent observations for landrace-fixed protein-coding variants using a strict fixation filter. A variant was classified as landrace-fixed only if the alternate allele was present in ≥8 of 20 tetraploid alleles (≥4 of 5 accessions) in one landrace and ≤2 of 20 in the other. This threshold excludes single-accession heterozygotes and segregating polymorphisms that do not represent landrace-level divergence. Sensitivity analysis confirmed that the fixation call was robust across thresholds: 245 variants at ≥8 alleles, 76 at ≥10, and 8 at ≥20 (full fixation). The headline variant survived at near-full-fixation thresholds (≥16 alleles) (Figure 5). The p.Trp384* locus at LT934119.1:416,494,333 is multi-allelic: three Mahmoudi accessions (SRR37252001, SRR37252002, SRR37252004) are homozygous for a simple SNV (C→T), and one (SRR37252003) is homozygous for a complex multinucleotide allele at the same site, both annotated as p.Trp384* by SnpEff; combined alternate-allele frequency = 0.80.

Of the 31 candidates with nominally HIGH-impact SnpEff calls, only one gene met the strict fixation criterion: TRITD_5Av1G148980, a TIR-NBS-LRR disease-resistance gene (Mahmoudi-elevated, ashr log□FC = −1.18, baseMean = 764; cis-co-localised with the chromosome 5A FST hotspot at FST = 0.830, ∼416.5 Mb on LT934119.1). All five Chili accessions were homozygous reference (0/0/0/0; AF = 0.00). Four of five Mahmoudi accessions were independently homozygous for a premature stop codon (1/1/1/1; AF = 0.80), with one Mahmoudi accession homozygous reference (AF = 0.00) (Figure 5e). The variant (LT934119.1:416,494,333 C→T; SnpEff: p.Trp384*, HIGH impact, stop_gained) truncates the 1,224-amino-acid protein at residue 383, eliminating the NB-ARC ATPase domain (residues ∼200–500), the LRR pathogen-recognition surface (residues ∼500–1,100), and the C-terminal tail. The retained N-terminal fragment comprises the TIR signalling domain (residues 1–∼160), a short linker (residues ∼160–200), and the N-terminal portion of the NB-ARC ATPase domain (residues ∼200–383). The C-terminal portion of NB-ARC (residues ∼384-500) is eliminated by the truncation. AlphaFold 3 prediction of the Svevo reference allele (mean pLDDT 67.3; TIR domain 67.8; NB-ARC 83.1; LRR 65.9; pTM 0.66) provided the structural template for annotating the truncation boundary (Figure 5a, b). A scan of the ±5 kb window surrounding TRITD_5Av1G148980 identified three landrace-fixed variants, all located in the coding sequence (including p.Trp384*); no variants passed quality filtering in the flanking non-coding regulatory regions (upstream, UTR, intronic, downstream).

Per-sample RNA-seq read counts confirmed that the Mahmoudi-truncated allele is not degraded by nonsense-mediated decay. Expression of TRITD_5Av1G148980 was 3.7-fold elevated in Mahmoudi (root: 1,746 reads; leaf: 892 reads) relative to Chili (root: 411 reads; leaf: 310 reads) (Figure 5c), consistent across both tissues (root VST log□FC = −2.16; leaf VST log□FC = −1.22; ashr log□FC = −1.18, p = 4.0 × 10□□). The data are therefore consistent with active transcription and likely translation of a TIR-domain-retaining protein fragment in Mahmoudi, not with NMD-mediated mRNA degradation. Coupled with the fixed coding truncation at p.Trp384*, this pattern is consistent with positive selection for retention of the TIR signalling module while eliminating the LRR pathogen-recognition and NB-ARC ATPase domains, aligning with the documented role of TIR-only NLR fragments as helper or regulatory decoys in plant immunity ( Lapin et al. 2019; Huang et al. 2022; Karasov et al. 2014). qRT-PCR with allele-specific primers and ribosome profiling will be required to formally separate transcriptional regulation from NMD efficiency.

The second cis-co-localised gene, TRITD_5Av1G148950 (chloroplast-movement protein, Chili-elevated, ashr log□FC = +1.34, baseMean = 79; same chromosome 5A FST hotspot), carried no landrace-fixed coding substitutions (Figure 5d). Two segregating missense variants were detected in the coding sequence (p.Glu653Gly and p.Cys554Ser), but both were heterozygous in a single Chili accession (AF_Chili = 0.25, AF_Mahmoudi = 0.00) and failed the strict fixation threshold. A scan of the ±5 kb cis-regulatory window identified no landrace-biased high-confidence SNPs (max |AF_diff| = 0.05). Per-sample RNA-seq read counts confirmed Chili-elevated expression (Chili root: 46 reads; Chili leaf: 141 reads; Mahmoudi root: 5 reads; Mahmoudi leaf: 42 reads; Chili/Mahmoudi ratio = 4.0×). Neither proximal coding nor proximal regulatory variation can therefore account for the observed Chili-elevated expression of this gene, indicating that the regulatory mechanism must lie outside the ±5 kb window, in distal cis-elements, trans-acting factors, or epigenetic differences not captured by short-read SNP calling.

Among the remaining 404 trans-regulated expression-divergent observations, 29 candidates carried nominally HIGH-impact SnpEff calls. After strict fixation filtering, zero landrace-fixed coding substitutions were identified. This structural-genomic negative result is internally consistent with the main paper’s conclusion that expression divergence in Tunisian durum wheat operates primarily through trans-regulatory rewiring rather than protein-coding change.

## 4. DISCUSSION

This study provides the first integrative whole-genome and whole-transcriptome characterisation of adaptive divergence between two Tunisian durum wheat landraces, Chili and Mahmoudi, cultivated under contrasting agro-ecological conditions. The multi-omic analysis uncovers a fundamental molecular architecture of local adaptation. Directional genomic selection and transcriptome regulatory divergence are spatially decoupled yet functionally convergent, challenging a purely cis-regulatory model of adaptive divergence (McManus et al., 2010; Wittkopp et al., 2004) and highlighting trans-regulatory evolution as a major mechanism shaping adaptive phenotypes in polyploid cereals.

The two landraces evolved under contrasting agro-ecological conditions: Chili mainly in central and northern Tunisia (Ben Krima et al., 2020), and Mahmoudi in the semi-arid south (Ouaja et al., 2021). This environmental contrast provides the ecological framework within which all subsequent genomic and transcriptomic signals must be interpreted, and constitutes the primary selective backdrop driving the divergence documented here.

Population structure analysis confirmed that Chili and Mahmoudi form genetically distinct clusters prior to any selection-based filtering, a prerequisite for the FST outlier framework: differentiation signals can only be interpreted as adaptive if the two groups are demonstrably non-panmictic at the genome level. The forty-six FST outlier windows distributed across six chromosomes collectively define a genomic architecture of discrete adaptive loci superimposed on a shared demographic background. This interpretation is supported by neutral marker statistics: Tajima’s D was uniformly negative in both landraces (median = −1.11), and nucleotide diversity was broadly comparable (Chili: 1.00 × 10□□; Mahmoudi: 1.11 × 10□□), consistent with a shared post-domestication expansion rather than population-specific bottlenecks. Genome-wide mean weighted FST was 0.083, confirming that background differentiation is modest and making the outlier peaks, which reach FST > 0.8 on chromosomes 5A and 6B, all the more striking by comparison. Per-chromosome nucleotide diversity revealed that Chili π exceeded Mahmoudi π specifically on chromosomes 5B, 6B, and 7B, a pattern consistent with selection having reduced diversity in Mahmoudi at these loci while Chili retained ancestral variation, further corroborating the directional selection model at the 6B hotspot in particular. Because the permutation null accounts for this shared demographic background, the discrete FST peaks cannot be attributed to genome-wide genetic drift (Beaumont and Nichols, 1996; Nielsen, 2005; Tajima, 1989). They point, instead, to real directional selection at specific adaptive loci.

Genome-wide scans across diverse tetraploid wheat accessions have repeatedly revealed signatures of selection concentrated on chromosomes 1A, 1B, 2A, 2B, 5A, and 6B, often co-localising with known QTL regions. Chromosomes 1A and 2A carry well-documented loci for drought tolerance and yield components, with chromosome 2A playing a particularly critical role through its influence on physiological and agronomical traits under water-limited conditions (Hu et al., 2020; Pshenichnikova et al., 2021). Chromosomes 1B and 2B harbour both drought-related yield QTLs and well-characterised disease resistance gene clusters, including loci for seedling drought tolerance and Fusarium head blight resistance (Arruda et al., 2016; Rabbi et al., 2021; Sari et al., 2018). Chromosome 5A harbours a strong FST signal in our dataset (FST = 0.830; empirical p = 0.013), spanning 10 outlier windows, making it one of the most prominent signatures of selection identified. This region coincides with a genomic interval known to host major phenological regulators, notably the vernalization gene VRN-A1, whose pleiotropic effects on flowering time and grain yield contribute to drought escape responses (Dolferus et al., 2019; Yan et al., 2003). Beyond VRN-A1, this signal includes two cis-co-localised expression candidates: a chloroplast movement gene (TRITD_5Av1G148950), elevated in Chili, and a TIR-NBS-LRR resistance gene (TRITD_5Av1G148980), elevated in Mahmoudi, suggesting that selective pressure on this chromosomal region simultaneously targets phenological adaptation and disease resistance modules. The cis-co-localised interval on chromosome 5A (∼416.5 Mb) exemplifies two distinct molecular mechanisms at a single selection hotspot. TRITD_5Av1G148980 (TIR-NBS-LRR, Mahmoudi-elevated) carries a near-fixed premature stop codon (p.Trp384*; AF_Mahmoudi = 0.80) that truncates the protein to a TIR-domain-retaining fragment, eliminating the C-terminal portion of the NB-ARC ATPase domain and the entire pathogen-recognition LRR domain. TIR-only NLR fragments are documented in plant immunity as helper or regulatory decoys that engage downstream EDS1/PAD4 signalling nodes to regulate defense, often acting as modules that retain enzymatic or signaling capacity without requiring canonical full-length activation (Lapin et al., 2019; Li & Tao, 2022; van Wersch et al., 2020). The 3.7-fold elevated expression of the truncated allele in Mahmoudi, despite the premature stop codon, indicates that the mRNA escapes nonsense-mediated decay and is actively transcribed, consistent with selection for retention of the TIR signalling module. In the lower-pathogen-pressure arid environment of Mahmoudi cultivation, selection against the severe energetic costs and autoimmunity risks traditionally associated with full-length NLRs (Karasov et al., 2014; Richard et al., 2018), while potentially retaining TIR-dependent NADase activity to prime basal defenses (Huang et al., 2022), represents a highly plausible adaptive trajectory. qRT-PCR with allele-specific primers and ribosome profiling will be required to formally separate transcriptional regulation from NMD efficiency. By contrast, TRITD_5Av1G148950 (chloroplast movement, Chili-elevated) shows no proximal coding or regulatory sequence divergence, indicating that its expression change is driven by distal or trans-acting mechanisms. The coexistence of these two modes, coding truncation in one gene, regulatory exclusion in the other, at the same chromosomal selection hotspot demonstrates that cis-co-localisation alone is insufficient to infer molecular mechanism, and that integrated structural-genomic and regulatory scanning is required to resolve adaptive architecture at individual loci.

Within this framework, chromosome 6B stands out as the most compelling signal in our dataset. The chromosome 6B hotspot (FST = 0.833; empirical p = 0.013) represents the strongest case for directional selection in this germplasm. Near-fixation levels of differentiation between closely related landraces are rare and suggest strong, sustained selective pressure concentrated on a small genomic region (Excoffier et al., 2009; Lewontin and Krakauer, 1973). Chromosome 6B is recognised more broadly as a principal carrier of major drought tolerance loci across diverse wheat germplasm, associated with grain yield stability under drought and combined heat and drought stress (Cavanagh et al., 2013; Nagy et al., 2022; Rabbi et al., 2021). Notably, its short arm includes the GPC-B1 (NAM-B1) locus, encoding a NAC-domain transcription factor involved in flag leaf senescence and the remobilization of nitrogen to the grain (Andleeb et al., 2023; Distelfeld et al., 2004; Uauy et al., 2006). Because this gene is closely linked to the transition from vegetative growth to grain filling, it directly affects nitrogen-use efficiency and grain protein content. Variation at this locus may therefore contribute to adaptation to contrasting water availability conditions, providing a mechanistically grounded explanation for the strong FST signal detected in this region.

A closer examination of chromosome 6B, however, reveals a complete absence of any relationship between the selection signal and local expression divergence. The 53 expression-divergent features mapped to this chromosome are genetically independent of the identified selection hotspot (r² ≤ 0.25 across all distances; physical separation > 1 Mb). The most parsimonious interpretation is that selection has acted on a coding or structural variant at approximately 154.4 Mb, most plausibly GPC-B1/NAM-B1 itself or a tightly linked regulatory element, while trans-acting regulators located elsewhere in the genome coordinate the expression programme of genes distributed across chromosome 6B.

This mechanistic decoupling is consistent with the known function of NAM-B1 as a transcriptional activator of senescence and nutrient remobilisation programmes. Divergence at this master regulator would propagate widespread expression changes across its downstream target network without requiring sequence variation in any individual target gene. The functional enrichment of the 53 expression-divergent features in disease resistance and stress-response categories adds another layer of support to this picture: genomic and transcriptomic signals are converging on the same adaptive pathways, but through distinct molecular mechanisms. This is a pattern of decoupled yet functionally coherent adaptation.

The predominance of trans-regulatory divergence observed genome-wide (99.5% at ±50 kb; 98.5% at ±500 kb) elevates this from a chromosome 6B-specific observation to a general architectural principle. Adaptation in these landraces operates not by evolving regulatory sequences adjacent to individual target genes, but by remodelling master hubs that coordinate entire adaptive gene batteries from remote chromosomal positions. This extends the King–Wilson regulatory hypothesis (King and Wilson, 1975) into the domain of quantitative genome-wide selection, consistent with the broader eQTL literature where trans-effects consistently predominate under adaptive divergence (Rachel B. Brem, 2001; Rockman and Kruglyak, 2006; Stranger et al., 2007), and with evidence from plant systems that populations adapted to contrasting climatic conditions evolve distinct transcriptome programmes constituting adaptive expression syndromes (Lasky et al., 2014; Zhang et al., 2016)

The functional coherence of this architecture is striking: trans-regulated observations are significantly enriched for disease resistance and protein homeostasis genes, precisely the categories under positive genomic selection, implying that selection targeted a small number of hubs with broad downstream reach. Disease resistance genes (NBS-LRR, RLK, and PR families) show the strongest enrichment signal in the dataset, consistent with evidence that trans-eQTL hotspots in cereals preferentially regulate immune gene networks from remote chromosomal positions (Abraham and Croll, 2023), while ubiquitin–proteasome pathway components constitute a second significantly enriched category, consistent with the proteostatic hub argument and the established role of the ubiquitin–26S proteasome system as a master transcriptional coordinator of plant stress and immune responses (Adams and Spoel, 2018; Vierstra, 2009). Together, these two categories demonstrate that the trans-regulatory architecture is not functionally random but is precisely calibrated on the adaptive modules that also bear the strongest genomic selection signatures.

WRKY and MYB transcription factors (25 of 28 TF observations Chili-elevated) are strong hub candidates (Eulgem et al., 2000; Pandey and Somssich, 2009; Ye et al., 2021), as are ubiquitin-proteasome components, whose divergence would reconfigure the proteostatic landscape of stress signalling through differential protein turnover rather than differential transcription of individual effector genes (Finley, 2009; Lyzenga and Stone, 2012; Smalle and Vierstra, 2004; Vierstra, 2009).

In polyploid wheat, sub-genome redundancy further facilitates this strategy by relaxing pleiotropic constraints: one homeologous copy can evolve novel trans-regulatory interactions while its partner maintains ancestral function (Adams and Wendel, 2005; Comai, 2005; Doyle et al., 2008; Feldman and Levy, 2009; Zhao et al., 2020). This provides an expanded regulatory search space in which hub rewiring becomes an evolutionarily accessible route to adaptive diversification.

This type of trans-regulatory organisation does not appear to be unique to the landraces studied here. In hexaploid wheat, the transcription factor TaMYB7-A1 acts as a central regulator, coordinating the expression of numerous drought-responsive genes through trans-eQTL interactions. This is corroborated at much larger scale by a multi-omics study combining GWAS, population-level transcriptomics across 228 wheat accessions, and eQTL mapping on a core set of 110 genotypes, which demonstrated that trans-acting regulatory variants dominate the control of gene expression under drought stress conditions, with condition-specific eQTL hotspots significantly enriched in pathways linked to water-use efficiency, including ABA signalling, stomatal movement regulation, and water deprivation response (Zhou et al., 2025). Taken together, these findings support the conclusion that adaptation in wheat is primarily mediated by key regulatory master hubs rather than by the independent selection of individual downstream target genes.

The contrasting water-economy strategies of the two landraces translate the genomic signals into physiological logic. Chili’s elevated aquaporins (PIP and TIP subfamily members) support an “acquire-and-distribute” strategy suited to adequate water supply (Chaumont and Tyerman, 2014; Maurel et al., 2015), maximising symplastic water flux through roots and source-to-sink distribution through leaves. Overexpression of the durum wheat aquaporin TdPIP2;1 has been shown to confer drought and salt tolerance in transgenic lines (Ayadi et al., 2019; Ayadi et al., 2011), validating the functional significance of natural variation in this gene family. Mahmoudi’s constitutive upregulation of dehydrins, LEA proteins, and ROS-scavenging enzymes, including superoxide dismutases, catalases, and peroxidase, reflects a “store-and-protect” programme characteristic of semi-arid adaptation (Close, 1996; Miller et al., 2010; Mittler, 2002; Tunnacliffe and Wise, 2007).

Dehydrins and LEA proteins are well known for their role as molecular chaperones, stabilising membranes and proteins during cellular dehydration (Kovacs et al., 2008). Their constitutive expression likely provides a first layer of protection against the rapid onset of water deficit, characteristic of the semi-arid south (Liu, Hao et al., 2020a).

Beyond these water-economy mechanisms, the higher abundance of seedling storage proteins in Chili may indicate a greater investment in nitrogen reserves. Such reserves are essential to sustain early growth under more favourable, mesic conditions, in agreement with the established role of storage proteins in fuelling germination and early seedling development (Cao et al., 2022; Shewry and Halford, 2002; Wen et al., 2018).

Chaperone enrichment in Mahmoudi, meanwhile, suggests that this landrace appears to keep its protective machinery running as a baseline. Its chaperone proteins act as a standing defence that starts before damage occurs, holding cellular order together as temperatures climb to protein-denaturing levels. This preparation makes biological sense for a landrace shaped by the intense, recurring heat of southern Tunisia. It also mirrors what has been observed in durum wheat landraces from similarly demanding environments, where heat shock proteins function as constitutive molecular chaperones, stabilising proteins and preserving cellular homeostasis under high-temperature conditions (De Pascali et al., 2022; Rampino et al., 2009). The same constitutive logic extends to oxidative stress management. The strong Mahmoudi bias of ROS-scavenging genes (96.9%; 31 of 32 manually curated observations) deviates significantly from a 1:1 null expectation (one-tailed binomial test, p ≈ 1.5 × 10□□), supporting adaptive rather than neutral divergence: random regulatory drift would be expected to produce a roughly equal bidirectional distribution, analogous to the 1:1 ratio observed for disease resistance genes. This asymmetry aligns with constitutive antioxidant priming. This physiological state, reported in drought-adapted varieties, allows faster clearance of ROS when stress hits (Conrath et al., 2006; Mittler, 2002). Drought priming research in wheat gives solid backing to this idea, showing that early exposure to moderate water deficit leads to higher antioxidant enzyme activity and better leaf water retention under subsequent stress (Amini et al., 2023). That functional outcome is precisely what the priming model predicts and provides a proof of concept that Mahmoudi’s constitutive antioxidant profile may represent a plausible source of adaptive advantage. However, it remains to be confirmed through direct stress-challenge experiments, required by the priming hypothesis (Beckers and Conrath, 2007; Conrath et al., 2006), in order to confirm that a pre-activated physiological state translates into measurable resilience, a dimension that constitutive seedling transcriptomes alone cannot capture.

Beyond the interpretations above, cross-study validation supports the biological relevance of the observed expression divergence. Matching against published wheat stress transcriptomes (Aprile et al., 2009; Habash et al., 2014; Lee et al., 2022; Liu, Haipei et al., 2020b) reveals a dual profile.

Overlaps with known stress genes show the identified seedling divergences match adaptive traits reported elsewhere. However, the majority fraction reflect novel germplasm-specific regulatory patterns absent from existing wheat stress datasets (Zuluaga et al., 2023). This novel fraction underscores the value of characterising traditional North African landraces, whose regulatory architectures remain largely unexplored relative to elite germplasm and may represent untapped genetic diversity for breeding.

From a breeding perspective, these results are particularly relevant for wheat improvement in North Africa, where enhancing drought resilience remains a key objective. The chromosomal hotspots identified here offer molecularly anchored targets suitable for marker-assisted selection, including cis-co-localised resistance gene candidates of direct breeding relevance given the sustained demand for novel rust resistance alleles in durum wheat (Luo et al., 2022; Periyannan et al., 2013; Singh et al., 2011).

Genomics-driven approaches integrating population genomics with participatory varietal selection have demonstrated their potential for breeding locally adapted varieties in smallholder systems (Albert and Kruglyak, 2015; Rockman and Kruglyak, 2006), and the hotspot regions identified here provide concrete molecular anchors for such programmes. However, the near-complete trans-regulatory architecture means that key adaptive expression phenotypes, including elevated disease resistance gene expression, ROS scavenging, and dehydrin accumulation, cannot be captured by introgression of single chromosomal intervals, because the regulatory variants controlling them map to remote positions elsewhere in the genome. Trans-regulated adaptive traits require eQTL mapping in segregating populations to identify causal regulatory variants before marker-assisted pyramiding becomes feasible (Albert and Kruglyak, 2015; Rockman and Kruglyak, 2006). This exemplifies a broader principle: the regulatory architecture of locally adapted landraces is a hidden layer of diversity that population genomics alone cannot access, and joint genomic–transcriptomic frameworks of the type demonstrated here are essential for its exploitation (McManus et al., 2010; Nica and Dermitzakis, 2013).

## 5. LIMITATIONS AND FUTURE DIRECTIONS

Several limitations temper these conclusions. RNA-seq replication of n = 1 per group precludes formal statistical inference; the 406 expression observations are exploratory and require biological replication (n ≥ 3) with qRT-PCR validation before confirmation (Conesa et al., 2016; Schurch et al., 2016). The WGS panel of n = 5 accessions per landrace is sufficient to identify extreme FST outliers but underpowered for population-level allele-frequency estimation and selection-scan methods that perform best at n ≥ 20 (Fumagalli et al., 2013; Gossmann et al., 2012; Hoban et al., 2016; Lotterhos and Whitlock, 2014). Tetraploid-to-diploid genotype conversion preserves information exactly for the 88.1% of homozygous calls but introduces potential dosage ambiguity for the 11.9% heterozygous fraction; across 277,770 genotype calls spanning 27,777 SNPs and 10 accessions, the dosage-class breakdown of this fraction was 7.9% one-allele heterozygotes, 2.7% two-allele, and 1.3% three-allele calls, meaning that exact genotype information was preserved for the 88.1% fully homozygous sites and potential dosage ambiguity is strictly limited to the heterozygous 11.9%; tetraploid-aware variant-calling tools should replace this conversion in subsequent analyses (Garrison and Marth, 2012; Serang et al., 2012). Chromosome-level FST × DEG co-localisation is conservative; bedtools gene-body intersection is recommended as a priority next analytical step. Functional enrichment relied on AHRD keyword matching rather than formal GO or KEGG annotation, and results should be treated as directional pending InterProScan and eggNOG annotation of the Svevo v1.0 reference (Maccaferri et al., 2019). Finally, all transcriptome data represent constitutive seedling conditions and cannot resolve dynamic stress response differences between the two landraces.

Future work will address these limitations in sequence: WGS expansion to ≥ 20 accessions per landrace to sharpen selection-scan power; RNA-seq replication with n ≥ 3 per group and tissue; stress-induction kinetics experiments under controlled drought; eQTL mapping in a Chili × Mahmoudi recombinant population to identify causal trans-regulatory variants; fine-mapping and allele-specific expression analysis at the chromosome 6B hotspot; and field trials under contrasting rainfall regimes across northern and southern Tunisia to connect molecular divergence with agronomic performance.

## 6. CONCLUSIONS

Taken together, our results establish that local adaptation in Tunisian durum wheat operates through a two-tier molecular architecture: discrete chromosomal islands of directional selection, and a genome-wide network of trans-regulatory divergence that targets the same functional modules, disease resistance, protein homeostasis, and oxidative stress management, via independent molecular mechanisms. The dominant chromosome 6B hotspot (FST = 0.833) exemplifies this architecture: strong selection at a coding or structural variant, with the broader chromosomal expression programme coordinated by remote trans-acting regulators. The two contrasting water-economy strategies, Chili’s “acquire-and-distribute” aquaporin programme and Mahmoudi’s “store-and-protect” dehydrin and antioxidant priming state, illustrate how this regulatory architecture translates genomic divergence into coherent and environmentally matched physiological phenotypes. This decoupled but functionally coherent multi-omic landscape, characterised here for the first time in North African wheat germplasm, provides both a conceptual framework for understanding cereal adaptation to Mediterranean climate gradients and a practical roadmap for exploiting the rich regulatory diversity of traditional landraces in next-generation breeding programmes.

### Data Availability

All raw sequencing data generated and analysed in this study are deposited at the NCBI Sequence Read Archive (SRA) under BioProject PRJNA1420514.

□ Whole-genome sequencing (WGS), Chili (n = 5): SRR37204970 – SRR37204974
□ Whole-genome sequencing (WGS), Mahmoudi (n = 5): SRR37252001 – SRR37252005
□ RNA-seq, Chili root and leaf: SRR37213950, SRR37213951
□ RNA-seq, Mahmoudi root and leaf: SRR37252182, SRR37252142

The Triticum turgidum cv. Svevo v1.0 reference genome and EMBL GFF3 gene models were obtained from NCBI (assembly accession GCA_900231445.1; chromosomes 1A-7B, accessions LT934111.1-LT934124.1).

All analysis scripts (WGS pipeline, RNA-seq pipeline, FST permutation testing, FST × DEG co-localisation, AHRD-keyword Fisher enrichment, sensitivity analyses, dosage cross-checks), processed data tables, the empirical permutation null distribution (perm_max_fst.txt, n = 1,000), and the interactive Plotly.js webapp are openly available at: https://github.com/nour0810/TuniTritica

The interactive webapp is also deployed via GitHub Pages at: https://nour0810.github.io/TuniTritica/

No proprietary or restricted-access data were used.

### Supplementary Data

The following files accompany the manuscript and are bundled with the GitHub repository (folder supplementary/):

#### Supplementary Tables

Table S1 – Table_S1_Landrace_Observations.csv. Full list of the 406 expression-divergent observations between Chili and Mahmoudi with shrunken log□fold change, ranked DESeq2 p-values, AHRD product description, and curated functional category.

Table S2 – Table_S2_Tissue_Candidates.csv. Complete tissue contrast (root vs. leaf) DEG results from the additive DESeq2 model.

Table S3 – Table_S3_Per_Chromosome_Genomics.csv. Per-chromosome counts of FST outlier windows, median π, and median Tajima’s D for each landrace.

Table S4 – Table_S4_Alignment_Statistics.csv. Per-library RNA-seq read counts, alignment rates, concordant unique alignment, and featureCounts assignment rates.

Table S5 – Table_S5_CrossRef_Validation.csv. Cross-study validation of the 406 observations against nine published wheat stress studies.

Table S5b – Table_S5_Deduplicated_Genes.txt. The 160 unique cross-referenced genes after deduplication across categories.

Table S6 – Table_S6_FST_DEG_Overlap.csv. The 46 permutation-significant FST outlier windows (10-kb analysis), with co-localised genes at ±50 kb and ±500 kb extensions.

Table S7 – Table_S7_CrossTissue_Consistent_Observations.csv. VST-normalised expression values per tissue, used to verify cross-tissue consistency under the additive model.

Table S8 – Table_S8_TransGene_Enrichment.csv. Fisher’s exact test results (one-tailed; Benjamini–Hochberg FDR) for the 12 enrichment-tested AHRD-keyword categories on the 404 trans-regulatory observations.

Table S9 – Table_S9_PerSample_Variant_Counts.csv. Per-sample WGS variant counts (range 1,532–4,313; median 2,232; outlier Chili accession SRR37204972 = 16,603).

Supplementary Table S10. Variant inventory for TRITD_5Av1G148980 and TRITD_5Av1G148950. Columns: gene_id, position, ref, alt, HGVS.p, SnpEff_impact, effect_type, AF_Chili, AF_Mahmoudi, AF_diff, strict_fixation_status.

Supplementary Table S11. Per-accession genotypes at p.Trp384* (LT934119.1:416,494,333). Columns: sample, landrace, tetraploid_GT, homozygosity_status, notes. Rows: 10 accessions (SRR37204970–SRR37204974, SRR37252001–SRR37252005). Note: The locus is multi-allelic, three Mahmoudi accessions (SRR37252001, 02, 04) are homozygous for a simple SNV (C→T) and one (SRR37252003) is homozygous for a complex multinucleotide allele at the same site, both annotated as p.Trp384* by SnpEff; combined alternate-allele frequency = 0.80.

Supplementary Table S12. AlphaFold 3 structural confidence metrics for TRITD_5Av1G148980 Svevo reference allele. Columns: residue, amino_acid, domain, pLDDT. Summary: domain-mean pLDDT (TIR, NB-ARC, LRR, C-terminal), overall mean pLDDT (67.3), pTM (0.66).

#### Supplementary Figures

Figure S1 – Figure_S1_Dispersion_Plot.svg. DESeq2 mean–dispersion fit confirming that the global trend is well estimated despite n = 1 per landrace × tissue.

Figure S2 – Figure_S2_Permuted_FST_Distribution.pdf (and .png). Empirical null distribution of the maximum FST under 1,000 label-shuffle permutations; observed maximum FST = 0.833 lies at the 99th percentile (empirical P = 0.013 under the (k + 1)/(N + 1) formula). Summary statistics in S2_statistics.txt.

Figure S3 – Figure_S3_Enrichment_OR_Plot.pdf (and .png). Odds-ratio forest plot of the 12 AHRD-keyword categories tested for enrichment among trans-regulatory observations.

Figure S4 – Figure_S4_LD_Decay_6B.pdf (and .png). Pairwise r² between the 6B lead SNP and SNPs within ±5 Mb (n = 50), showing rapid LD decay (median r² ≤ 0.25; zero observations with r² > 0.2).

Figure S5 – Figure_S5_Constitutive transcriptome divergence between Tunisian durum wheat landraces.pdf.

#### Supplementary Notes and sensitivity files

Supplementary Note S1 – Supplementary_Note_S1.txt. Distance threshold sensitivity for FST × expression co-localisation (±50 kb, ±500 kb, ±1 Mb).

Supplementary Note S2 – Supplementary_Note_S2.txt. Methodology and results of the chromosome 6B linkage disequilibrium analysis.

enrichment_sensitivity.txt – three sensitivity scenarios for the trans-regulatory enrichment (alternative background; cis exclusion; 10-category collapsed classification).

dosage_distribution.tsv – tetraploid-to-diploid genotype-class breakdown across the 277,770 genotype calls (88.1 % homozygous, 11.9 % heterozygous).

S2_statistics.txt – summary statistics for the empirical FST null distribution (N = 1,000 permutations; observed max = 0.833; P95 = 0.500; P99 = 0.833; empirical P = 0.013).

## Acknowledgement

The authors thank the OpenDurumGPT consortium for making the whole-genome sequencing data of the Tunisian durum wheat landraces Chili and Mahmoudi freely available on Zenodo (zenodo.org/communities/opendurumgpt/records), and for their commitment to open science and the preservation of Tunisian agrobiodiversity.

## Consent for publication

Not applicable.

## Competing interests

The authors declare that they have no competing interests.

## Funding

This research received no external funding.

## Authors’ contributions

M.G-B.A designed the study. N.E.H.M conducted the analysis. M.G-B.A, I.B and R.D contributed to data acquisition and analysis. N.E.H.M, M.G-B.A, I.B and R.D prepared and wrote the original draft.

## Notes

https://nour0810.github.io/TuniTritica/

https://github.com/nour0810/TuniTritica

### Competing Interest Statement

The authors have declared no competing interest.

### Summary of Updates

Applying a strict landrace-fixation filter (8 or more of 20 tetraploid alleles, with sensitivity at 10, 16, and 20) to all 406 expression-divergent observations revealed a single landrace-fixed protein-coding variant: a premature stop codon p.Trp384* in the TIR-NBS-LRR gene TRITD_5Av1G148980 (AF_Mahmoudi = 0.80; four of five Mahmoudi accessions homozygous; absent in all five Chili accessions). AlphaFold 3 prediction of the Svevo reference allele (mean pLDDT 67.3; pTM 0.66) provided the structural template, showing that the truncation retains an N-terminal fragment (TIR and N-terminal NB-ARC, residues 1 to 383) and eliminates the C-terminal NB-ARC ATPase region, the LRR pathogen-recognition surface, and the C-terminal tail. Symmetric 5 kb cis-regulatory scans of both chromosome 5A candidates uncovered two distinct molecular mechanisms at the same selection hotspot: coding truncation in TRITD_5Av1G148980 and complete proximal regulatory exclusion in the chloroplast-movement gene TRITD_5Av1G148950. Per-sample RNA-seq read counts (Mahmoudi root 1,746 vs. Chili root 411; 3.7-fold elevation) demonstrate that the truncated mRNA escapes nonsense-mediated decay and is actively transcribed, consistent with positive selection for retention of the TIR signalling module. Statistical support was added for the Mahmoudi ROS-bias asymmetry (one-tailed binomial, p approximately 1.5 x 10^-8). A new Methods Section 2.11, Figure 5, Supplementary Tables S10 to S12, and Notes S3 to S4 document these analyses. The TuniTritica webapp (https://nour0810.github.io/TuniTritica/) and reproducibility package (https://github.com/nour0810/TuniTritica) have been updated with all v3 scripts and outputs. Limitations, hedging, and exploratory framing remain unchanged.

## References

1. Abraham, L.N., Croll, D., 2023. Genome-wide expression QTL mapping reveals the highly dynamic regulatory landscape of a major wheat pathogen. BMC biology 21(1), 263.

2. Adams, E.H., Spoel, S.H., 2018. The ubiquitin–proteasome system as a transcriptional regulator of plant immunity. Journal of Experimental Botany 69(19), 4529–4537.

3. Adams, K.L., Wendel, J.F., 2005. Polyploidy and genome evolution in plants. Current opinion in plant biology 8(2), 135–141.

4. Albert, F.W., Kruglyak, L., 2015. The role of regulatory variation in complex traits and disease. Nature Reviews Genetics 16(4), 197–212.

5. Amini, A., Majidi, M.M., Mokhtari, N., Ghanavati, M., 2023. Drought stress memory in a germplasm of synthetic and common wheat: antioxidant system, physiological and morphological consequences. Scientific Reports 13(1), 8569.

6. Anderson, R., Bayer, P.E., Edwards, D., 2020. Climate change and the need for agricultural adaptation. Current opinion in plant biology 56, 197–202.

7. Andleeb, T., Knight, E., Borrill, P., 2023. Wheat NAM genes regulate the majority of early monocarpic senescence transcriptional changes including nitrogen remobilization genes. G3 13(2), jkac275.

8. Aprile, A., Mastrangelo, A.M., De Leonardis, A.M., Galiba, G., Roncaglia, E., Ferrari, F., De Bellis, L., Turchi, L., Giuliano, G., Cattivelli, L., 2009. Transcriptional profiling in response to terminal drought stress reveals differential responses along the wheat genome. BMC genomics 10(1), 279.

9. Arora, N.K., 2019. Impact of climate change on agriculture production and its sustainable solutions. Environmental sustainability 2(2), 95–96.

10. Arruda, M.P., Brown, P., Brown□Guedira, G., Krill, A.M., Thurber, C., Merrill, K.R., Foresman, B.J., Kolb, F.L., 2016. Genome□wide association mapping of Fusarium head blight resistance in wheat using genotyping□by□sequencing. The Plant Genome 9(1), plantgenome2015.2004.0028.

11. Ayadi, M., Brini, F., Masmoudi, K., 2019. Overexpression of a wheat aquaporin gene, Td PIP2; 1, enhances salt and drought tolerance in transgenic durum wheat cv. Maali. International Journal of Molecular Sciences 20(10), 2389.

12. Ayadi, M., Cavez, D., Miled, N., Chaumont, F., Masmoudi, K., 2011. Identification and characterization of two plasma membrane aquaporins in durum wheat (Triticum turgidum L. subsp. durum) and their role in abiotic stress tolerance. Plant Physiology and Biochemistry 49(9), 1029–1039.

13. Ayed, S., Babay, E., Ben Amor, A., Ben M’Barek, S., Djemal, R., Ebel, C., Elleuch, A., Gargouri, S., Ghanmi, S., Ghazala, I., Hamza, S., Hanin, M., Medini, M., Robbana, C., Saidi, M.N., Slim, A., Sayahi, N., 2026. From Local Heritage to Genomic Innovation: Whole-Genome Sequencing of Tunisian Durum Wheat Landraces Chili and Mahmoudi Zenodo. 10.5281/zenodo.19499032.

14. Beaumont, M.A., Nichols, R.A., 1996. Evaluating loci for use in the genetic analysis of population structure. Proceedings of the Royal Society of London. Series B: Biological Sciences 263(1377), 1619–1626.

15. Beckers, G.J., Conrath, U., 2007. Priming for stress resistance: from the lab to the field. Current opinion in plant biology 10(4), 425–431.

16. Ben Krima, S., Slim, A., Gélisse, S., Kouki, H., Nadaud, I., Sourdille, P., Yahyaoui, A., Ben M’barek, S., Suffert, F., Marcel, T.C., 2020. Life story of Tunisian durum wheat landraces revealed by their genetic and phenotypic diversity. bioRxiv, 2020.2008. 2014.251157.

17. Cao, H., Duncan, O., Millar, A.H., 2022. The molecular basis of cereal grain proteostasis. Essays in Biochemistry 66(2), 243–253.

18. Cavanagh, C.R., Chao, S., Wang, S., Huang, B.E., Stephen, S., Kiani, S., Forrest, K., Saintenac, C., Brown-Guedira, G.L., Akhunova, A., 2013. Genome-wide comparative diversity uncovers multiple targets of selection for improvement in hexaploid wheat landraces and cultivars. Proceedings of the national academy of sciences 110(20), 8057–8062.

19. Chaouachi, L., Marín-Sanz, M., Kthiri, Z., Boukef, S., Harbaoui, K., Barro, F., Karmous, C., 2023. The opportunity of using durum wheat landraces to tolerate drought stress: screening morpho-physiological components. AoB Plants 15(3), plad022.

20. Chaumont, F., Tyerman, S.D., 2014. Aquaporins: highly regulated channels controlling plant water relations. Plant physiology 164(4), 1600–1618.

21. Close, T.J., 1996. Dehydrins: emergence of a biochemical role of a family of plant dehydration proteins. Physiologia plantarum 97(4), 795–803.

22. Comai, L., 2005. The advantages and disadvantages of being polyploid. Nature reviews genetics 6(11), 836–846.

23. Conesa, A., Madrigal, P., Tarazona, S., Gomez-Cabrero, D., Cervera, A., McPherson, A., Szcześniak, M.W., Gaffney, D.J., Elo, L.L., Zhang, X., 2016. A survey of best practices for RNA-seq data analysis. Genome biology 17(1), 13.

24. Conrath, U., Beckers, G.J., Flors, V., García-Agustín, P., Jakab, G., Mauch, F., Newman, M.-A., Pieterse, C.M., Poinssot, B., Pozo, M.J., 2006. Priming: getting ready for battle. Molecular plant-microbe interactions 19(10), 1062–1071.

25. De Pascali, M., De Caroli, M., Aprile, A., Miceli, A., Perrotta, C., Gullì, M., Rampino, P., 2022. Drought stress pre-treatment triggers thermotolerance acquisition in durum wheat. International Journal of Molecular Sciences 23(14), 7988.

26. Distelfeld, A., Uauy, C., Olmos, S., Schlatter, A.R., Dubcovsky, J., Fahima, T., 2004. Microcolinearity between a 2-cM region encompassing the grain protein content locus Gpc-6B1 on wheat chromosome 6B and a 350-kb region on rice chromosome 2. Functional & Integrative Genomics 4(1), 59–66.

27. Dolferus, R., Thavamanikumar, S., Sangma, H., Kleven, S., Wallace, X., Forrest, K., Rebetzke, G., Hayden, M., Borg, L., Smith, A., 2019. Determining the genetic architecture of reproductive stage drought tolerance in wheat using a correlated trait and correlated marker effect model. G3: Genes, Genomes, Genetics 9(2), 473–489.

28. Doyle, J.J., Flagel, L.E., Paterson, A.H., Rapp, R.A., Soltis, D.E., Soltis, P.S., Wendel, J.F., 2008. Evolutionary genetics of genome merger and doubling in plants. Annual review of genetics 42(1), 443–461.

29. Eulgem, T., Rushton, P.J., Robatzek, S., Somssich, I.E., 2000. The WRKY superfamily of plant transcription factors. Trends in plant science 5(5), 199–206.

30. Excoffier, L., Hofer, T., Foll, M., 2009. Detecting loci under selection in a hierarchically structured population. Heredity 103(4), 285–298.

31. Faostat, F., 2023. Crops and livestock products. Food and Agriculture Organization of the United Nations.

32. Feldman, M., Levy, A.A., 2009. Genome evolution in allopolyploid wheat-a revolutionary reprogramming followed by gradual changes. Journal of Genetics and Genomics 36(9), 511–518.

33. Finley, D., 2009. Recognition and processing of ubiquitin-protein conjugates by the proteasome. Annual review of biochemistry 78(1), 477–513.

34. Fumagalli, M., Vieira, F.G., Korneliussen, T.S., Linderoth, T., Huerta-Sánchez, E., Albrechtsen, A., Nielsen, R., 2013. Quantifying population genetic differentiation from next-generation sequencing data. Genetics 195(3), 979–992.

35. Garrison, E., Marth, G., 2012. Haplotype-based variant detection from short-read sequencing. arXiv preprint arXiv:1207.3907.

36. Giorgi, F., Lionello, P., 2008. Climate change projections for the Mediterranean region. Global and planetary change 63(2-3), 90–104.

37. Gossmann, T.I., Keightley, P.D., Eyre-Walker, A., 2012. The effect of variation in the effective population size on the rate of adaptive molecular evolution in eukaryotes. Genome biology and evolution 4(5), 658–667.

38. Habash, D.Z., Baudo, M., Hindle, M., Powers, S.J., Defoin-Platel, M., Mitchell, R., Saqi, M., Rawlings, C., Latiri, K., Araus, J.L., 2014. Systems responses to progressive water stress in durum wheat. PloS one 9(9), e108431.

39. Huang, S., Jia, A., Song, W., Hessler, G., Meng, Y., Sun, Y., Xu, L., Laessle, H., Jirschitzka, J., Ma, S., Xiao, Y., Yu, D., Hou, J., Liu, R., Sun, H., Liu, X., Han, Z., Chang, J., Parker, J. E., & Chai, J. (2022). Identification and receptor mechanism of TIR-catalyzed small molecules in plant immunity. Science, 377(6605), eabq3297. 10.1126/science.abq3297

40. Hoban, S., Kelley, J.L., Lotterhos, K.E., Antolin, M.F., Bradburd, G., Lowry, D.B., Poss, M.L., Reed, L.K., Storfer, A., Whitlock, M.C., 2016. Finding the genomic basis of local adaptation: pitfalls, practical solutions, and future directions. The American Naturalist 188(4), 379–397.

41. Hu, J., Wang, X., Zhang, G., Jiang, P., Chen, W., Hao, Y., Ma, X., Xu, S., Jia, J., Kong, L., 2020. QTL mapping for yield-related traits in wheat based on four RIL populations. Theoretical and Applied Genetics 133(3), 917–933.

42. Karasov, T. L., Kniskern, J. M., Gao, L., DeYoung, B. J., Ding, J., Dubiella, U., Lastra, R. O., Nallu, S., Roux, F., Innes, R. W., Barrett, L. G., Hudson, R. R., & Bergelson, J. (2014). The long-term maintenance of a resistance polymorphism through diffuse interactions. Nature, 512(7515), 436–440. 10.1038/nature13439

43. King, M.-C., Wilson, A.C., 1975. Evolution at two levels in humans and chimpanzees: Their macromolecules are so alike that regulatory mutations may account for their biological differences. Science 188(4184), 107–116.

44. Kovacs, D., Kalmar, E., Torok, Z., Tompa, P., 2008. Chaperone activity of ERD10 and ERD14, two disordered stress-related plant proteins. Plant physiology 147(1), 381–390.

45. Lapin, D., Kovacova, V., Sun, X., Dongus, J., Bhandari, D. D., von Born, P., Bautor, J., Guarneri, N., Stuttmann, J., Beyer, A., & Parker, J. E. (2019). A coevolved EDS1-SAG101-NRG1 module mediates cell death signaling by TIR-domain immune receptors. The Plant Cell, 31(10), 2430–2448. 10.1101/572826

46. Lasky, J.R., Des Marais, D.L., Lowry, D.B., Povolotskaya, I., McKay, J.K., Richards, J.H., Keitt, T.H., Juenger, T.E., 2014. Natural variation in abiotic stress responsive gene expression and local adaptation to climate in Arabidopsis thaliana. Molecular biology and evolution 31(9), 2283–2296.

47. Latiri, K., Lhomme, J.-P., Annabi, M., Setter, T.L., 2010. Wheat production in Tunisia: Progress, inter-annual variability and relation to rainfall. European Journal of Agronomy 33(1), 33–42.

48. Lee, M.H., Kim, K.-M., Sang, W.-G., Kang, C.-S., Choi, C., 2022. Comparison of gene expression changes in three wheat varieties with different susceptibilities to heat stress using RNA-Seq analysis. International Journal of Molecular Sciences 23(18), 10734.

49. Lewontin, R.C., Krakauer, J., 1973. Distribution of gene frequency as a test of the theory of the selective neutrality of polymorphisms. Genetics 74(1), 175–195.

50. Li, J., & Tao, X. (2022). EDS1 modules as two-tiered receptor complexes for TIR-catalyzed signaling molecules to activate plant immunity. Stress Biology, 2, 1–4. 10.1007/s44154-022-00056-z

51. Liu, H., Yang, Y., Liu, D., Wang, X., Zhang, L., 2020a. Transcription factor TabHLH49 positively regulates dehydrin WZY2 gene expression and enhances drought stress tolerance in wheat. BMC plant biology 20(1), 259.

52. Liu, H., Able, A.J., Able, J.A., 2020b. Integrated analysis of small RNA, transcriptome, and degradome sequencing reveals the water-deficit and heat stress response network in durum wheat. International journal of molecular sciences 21(17), 6017.

53. Lopes, M.S., El-Basyoni, I., Baenziger, P.S., Singh, S., Royo, C., Ozbek, K., Aktas, H., Ozer, E., Ozdemir, F., Manickavelu, A., 2015. Exploiting genetic diversity from landraces in wheat breeding for adaptation to climate change. Journal of experimental botany 66(12), 3477–3486.

54. Lotterhos, K.E., Whitlock, M.C., 2014. Evaluation of demographic history and neutral parameterization on the performance of FST outlier tests. Molecular ecology 23(9), 2178–2192.

55. Luo, J., Rouse, M.N., Hua, L., Li, H., Li, B., Li, T., Zhang, W., Gao, C., Wang, Y., Dubcovsky, J., 2022. Identification and characterization of Sr22b, a new allele of the wheat stem rust resistance gene Sr22 effective against the Ug99 race group. Plant biotechnology journal 20(3), 554–563.

56. Lyzenga, W.J., Stone, S.L., 2012. Abiotic stress tolerance mediated by protein ubiquitination. Journal of experimental botany 63(2), 599–616.

57. Maccaferri, M., Harris, N.S., Twardziok, S.O., Pasam, R.K., Gundlach, H., Spannagl, M., Ormanbekova, D., Lux, T., Prade, V.M., Milner, S.G., 2019. Durum wheat genome highlights past domestication signatures and future improvement targets. Nature genetics 51(5), 885–895.

58. Maurel, C., Boursiac, Y., Luu, D.-T., Santoni, V., Shahzad, Z., Verdoucq, L., 2015. Aquaporins in plants. Physiological reviews.

59. McManus, C.J., Coolon, J.D., Duff, M.O., Eipper-Mains, J., Graveley, B.R., Wittkopp, P.J., 2010. Regulatory divergence in Drosophila revealed by mRNA-seq. Genome research 20(6), 816–825.

60. Meddi, M., Eslamian, S., 2021. Uncertainties in rainfall and water resources in Maghreb countries under climate change, African handbook of climate change adaptation. Springer, pp. 1967–2003.

61. Miazzi, M.M., Babay, E., De Vita, P., Montemurro, C., Chaabane, R., Taranto, F., Mangini, G., 2022. Comparative genetic analysis of durum wheat landraces and cultivars widespread in Tunisia. Frontiers in Plant Science 13, 939609.

62. Miller, G., Suzuki, N., Ciftci□Yilmaz, S., Mittler, R., 2010. Reactive oxygen species homeostasis and signalling during drought and salinity stresses. Plant, cell & environment 33(4), 453–467.

63. Mittler, R., 2002. Oxidative stress, antioxidants and stress tolerance. Trends in plant science 7(9), 405–410.

64. Nagy, É., Szabó-Hevér, Á., Lehoczki-Krsjak, S., Lantos, C., Kiss, E., Pauk, J., 2022. Detection of drought tolerance-related QTL in the Plainsman V./Cappelle Desprez doubled haploid wheat population. Cereal Research Communications 50(4), 689–698.

65. Nica, A.C., Dermitzakis, E.T., 2013. Expression quantitative trait loci: present and future. Philosophical Transactions of the Royal Society B: Biological Sciences 368(1620).

66. Nielsen, R., 2005. Molecular signatures of natural selection. Annu. Rev. Genet. 39(1), 197–218.

67. Ouaja, M., Bahri, B.A., Aouini, L., Ferjaoui, S., Medini, M., Marcel, T.C., Hamza, S., 2021. Morphological characterization and genetic diversity analysis of Tunisian durum wheat (Triticum turgidum var. durum) accessions. BMC Genomic Data 22(1), 3.

68. Pandey, S.P., Somssich, I.E., 2009. The role of WRKY transcription factors in plant immunity. Plant physiology 150(4), 1648–1655.

69. Periyannan, S., Moore, J., Ayliffe, M., Bansal, U., Wang, X., Huang, L., Deal, K., Luo, M., Kong, X., Bariana, H., 2013. The gene Sr33, an ortholog of barley Mla genes, encodes resistance to wheat stem rust race Ug99. Science 341(6147), 786–788.

70. Pshenichnikova, T.A., Osipova, S.V., Smirnova, O.G., Leonova, I.N., Permyakova, M.D., Permyakov, A.V., Rudikovskaya, E.G., Konstantinov, D.K., Verkhoturov, V.V., Lohwasser, U., 2021. Regions of chromosome 2A of bread wheat (Triticum aestivum L.) associated with variation in physiological and agronomical traits under contrasting water regimes. Plants 10(5), 1023.

71. Rabbi, S.H.A., Kumar, A., Mohajeri Naraghi, S., Simsek, S., Sapkota, S., Solanki, S., Alamri, M.S., Elias, E.M., Kianian, S., Missaoui, A., 2021. Genome-wide association mapping for yield and related traits under drought stressed and non-stressed environments in wheat. Frontiers in Genetics 12, 649988.

72. Rachel B. Brem, G.Y., Rebecca Clinton, Leonid Kruglyak, 2001. Genetic Dissection of Transcriptional Regulation in. Nature 411, 41.

73. Rampino, P., Mita, G., Pataleo, S., De Pascali, M., Di Fonzo, N., Perrotta, C., 2009. Acquisition of thermotolerance and HSP gene expression in durum wheat (Triticum durum Desf.) cultivars. Environmental and experimental Botany 66(2), 257–264.

74. Reynolds, M., Langridge, P., 2016. Physiological breeding. Current opinion in plant biology 31, 162–171.

75. Richard, M. M. S., Gratias, A., Meyers, B. C., & Geffroy, V. (2018). Molecular mechanisms that limit the costs of NLR□mediated resistance in plants. Molecular Plant Pathology, 19(11), 2516–2523. 10.1111/mpp.12723

76. Robbana, C., Kehel, Z., Ammar, K., Guzmán, C., Naceur, M.B.B., Amri, A., 2021. Unlocking the patterns of the Tunisian durum wheat landraces genetic structure based on phenotypic characterization in relation to farmer’s vernacular name. Agronomy 11(4), 634.

77. Robbana, C., Kehel, Z., Ben Naceur, M.b., Sansaloni, C., Bassi, F., Amri, A., 2019. Genome-wide genetic diversity and population structure of Tunisian durum wheat landraces based on DArTseq technology. International journal of molecular sciences 20(6), 1352.

78. Rockman, M.V., Kruglyak, L., 2006. Genetics of global gene expression. Nature Reviews Genetics 7(11), 862–872.

79. Royo, C., Maccaferri, M., Álvaro, F., Moragues, M., Sanguineti, M.C., Tuberosa, R., Maalouf, F., del Moral, L.F.G., Demontis, A., Rhouma, S., 2010. Understanding the relationships between genetic and phenotypic structures of a collection of elite durum wheat accessions. Field Crops Research 119(1), 91–105.

80. Sari, E., Berraies, S., Knox, R.E., Singh, A.K., Ruan, Y., Cuthbert, R.D., Pozniak, C.J., Henriquez, M.A., Kumar, S., Burt, A.J., 2018. High density genetic mapping of Fusarium head blight resistance QTL in tetraploid wheat. PloS one 13(10), e0204362.

81. Schlenker, W., Roberts, M.J., 2009. Nonlinear temperature effects indicate severe damages to US crop yields under climate change. Proceedings of the National Academy of sciences 106(37), 15594–15598.

82. Schurch, N.J., Schofield, P., Gierliński, M., Cole, C., Sherstnev, A., Singh, V., Wrobel, N., Gharbi, K., Simpson, G.G., Owen-Hughes, T., 2016. How many biological replicates are needed in an RNA-seq experiment and which differential expression tool should you use? Rna 22(6), 839–851.

83. Serang, O., Mollinari, M., Garcia, A.A.F., 2012. Efficient exact maximum a posteriori computation for Bayesian SNP genotyping in polyploids. PloS one 7(2), e30906.

84. Shewry, P.R., Halford, N.G., 2002. Cereal seed storage proteins: structures, properties and role in grain utilization. Journal of experimental botany 53(370), 947–958.

85. Singh, R.P., Hodson, D.P., Huerta-Espino, J., Jin, Y., Bhavani, S., Njau, P., Herrera-Foessel, S., Singh, P.K., Singh, S., Govindan, V., 2011. The emergence of Ug99 races of the stem rust fungus is a threat to world wheat production. Annual review of phytopathology 49(1), 465–481.

86. Smalle, J., Vierstra, R.D., 2004. The ubiquitin 26S proteasome proteolytic pathway. Annu. Rev. Plant Biol. 55(1), 555–590.

87. Stephens, M., 2017. False discovery rates: a new deal. Biostatistics 18(2), 275–294.

88. Stranger, B.E., Forrest, M.S., Dunning, M., Ingle, C.E., Beazley, C., Thorne, N., Redon, R., Bird, C.P., De Grassi, A., Lee, C., 2007. Relative impact of nucleotide and copy number variation on gene expression phenotypes. Science 315(5813), 848–853.

89. Tajima, F., 1989. Statistical method for testing the neutral mutation hypothesis by DNA polymorphism. Genetics 123(3), 585–595.

90. Tunnacliffe, A., Wise, M.J., 2007. The continuing conundrum of the LEA proteins. Naturwissenschaften 94(10), 791–812.

91. Uauy, C., Distelfeld, A., Fahima, T., Blechl, A., Dubcovsky, J., 2006. A NAC gene regulating senescence improves grain protein, zinc, and iron content in wheat. Science 314(5803), 1298–1301.

92. van Wersch, S., Tian, L., Hoy, R., & Li, X. (2020). Plant NLRs: The whistleblowers of plant immunity. Plant Communications, 1(1), 100090. 10.1016/j.xplc.2020.100090

93. Vierstra, R.D., 2009. The ubiquitin–26S proteasome system at the nexus of plant biology. Nature reviews Molecular cell biology 10(6), 385–397.

94. Weir, B.S., Cockerham, C.C., 1984. Estimating F-statistics for the analysis of population structure. evolution, 1358–1370.

95. Wen, D., Hou, H., Meng, A., Meng, J., Xie, L., Zhang, C., 2018. Rapid evaluation of seed vigor by the absolute content of protein in seed within the same crop. Scientific reports 8(1), 5569.

96. Whitehead, A., Crawford, D.L., 2006. Variation within and among species in gene expression: raw material for evolution. Molecular ecology 15(5), 1197–1211.

97. Wittkopp, P.J., Haerum, B.K., Clark, A.G., 2004. Evolutionary changes in cis and trans gene regulation. Nature 430(6995), 85–88.

98. Wray, G.A., 2007. The evolutionary significance of cis-regulatory mutations. Nature Reviews Genetics 8(3), 206–216.

99. Yan, L., Loukoianov, A., Tranquilli, G., Helguera, M., Fahima, T., Dubcovsky, J., 2003. Positional cloning of the wheat vernalization gene VRN1. Proceedings of the National Academy of Sciences 100(10), 6263–6268.

100. Ye, H., Qiao, L., Guo, H., Guo, L., Ren, F., Bai, J., Wang, Y., 2021. Genome-wide identification of wheat WRKY gene family reveals that TaWRKY75-A is referred to drought and salt resistances. Frontiers in Plant Science 12, 663118.

101. Zeven, A.C., 1998. Landraces: a review of definitions and classifications. Euphytica 104(2), 127–139.

102. Zhang, N., Vierling, E., Tonsor, S.J., 2016. Adaptive divergence in transcriptome response to heat and acclimation in Arabidopsis thaliana plants from contrasting climates. bioRxiv, 044446.

103. Zhao, N., Dong, Q., Nadon, B.D., Ding, X., Wang, X., Dong, Y., Liu, B., Jackson, S.A., Xu, C., 2020. Evolution of homeologous gene expression in polyploid wheat. Genes 11(12), 1401.

104. Zhou, Y., Wang, D., Wang, H., Qiao, Y., Zhao, P., Cao, Y., Liu, X., Yang, Y., Lin, X., Xu, S., 2025. Integrative omics of the genetic basis for wheat WUE and drought resilience reveal the function of TaMYB7-A1. Nature Communications 16(1), 8622.

105. Zuluaga, D.L., Blanco, E., Mangini, G., Sonnante, G., Curci, P.L., 2023. A survey of the transcriptomic resources in durum wheat: stress responses, data integration and exploitation. Plants 12(6), 1267.

