## Supplementary material for "Unlocking open-access genomic and transcriptomic data: the first bioinformatic exploitation of tunisian durum wheat landraces Chili and Mahmoudi": enrichment_sensitivity.txt

Enrichment sensitivity analyses — trans-regulatory observations
================================================================

Three sensitivity scenarios were run for the Fisher's exact test (one-tailed,
greater) on AHRD-keyword functional categories with Benjamini-Hochberg FDR
correction. Reference (primary) analysis: trans set = 404 observations,
background = 35,535 non-divergent expressed genes, 12 enrichment-tested
biological categories.

(a) Background = all 35,939 expressed genes (vs. 35,535 non-divergent)
    Disease resistance       : odds ratio = 2.6, FDR = 2.6e-9   [significant]
    Ubiquitin / proteasome   : odds ratio = 1.6, FDR = 0.054    [marginally non-significant]

(b) Trans set = 404 observations (2 cis genes excluded from the 406 total)
    Disease resistance       : odds ratio = 2.6, FDR = 1.4e-9   [significant; identical to primary]
    Ubiquitin / proteasome   : odds ratio = 1.6, FDR = 0.049    [significant; identical to primary]

(c) 10-category collapsed classification (vs. 12-category)
    Disease resistance       : odds ratio = 2.7, FDR = 5.8e-11  [strengthened]
    Ubiquitin / proteasome   : odds ratio = 1.6, FDR = 0.037    [significant]

Conclusion:
  Disease resistance enrichment is robust across all three sensitivity scenarios.
  Ubiquitin-proteasome enrichment is borderline (FDR 0.037-0.054) and is
  reported with the caveat that significance is scenario-dependent.

Script: scripts/04_enrichment/enrichment_sensitivity.py
