## Supplementary material for "Unlocking open-access genomic and transcriptomic data: the first bioinformatic exploitation of tunisian durum wheat landraces Chili and Mahmoudi": Figure_S5_Constitutive transcriptome divergence between Tunisian durum wheat landraces.pdf

a

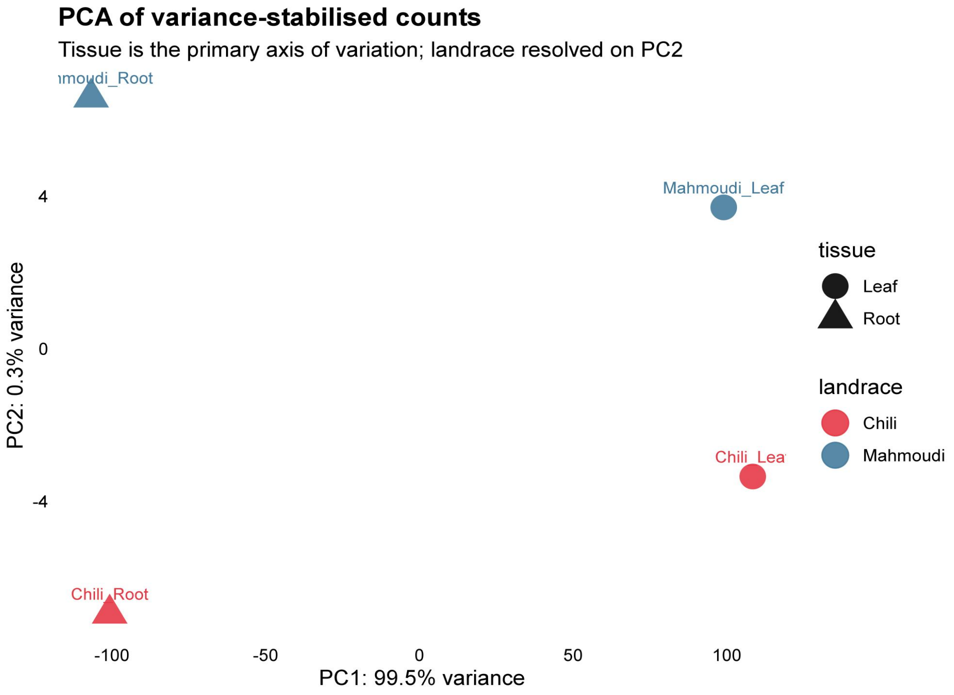
PCA of variance-stabilised counts
Tissue is the primary axis of variation; landrace resolved on PC2
ımoudi_Root
Mahmoudi_Leaf
Chili_Lea
Chili_Root
PC2: 0.3% variance
4
0
-4
-100
-50
0
50
100
PC1: 99.5% variance
tissue
Leaf
Root
landrace
Chili
Mahmoudi

b

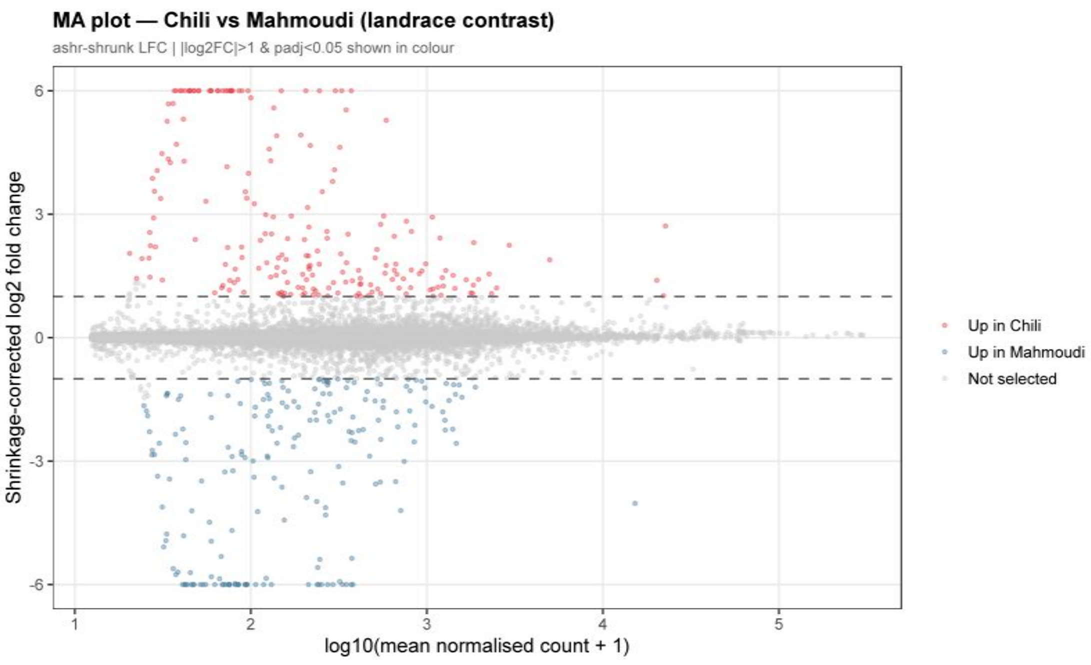
MA plot — Chili vs Mahmoudi (landrace contrast)
ashr-shrunk LFC | |log2FC|>1 & padj<0.05 shown in colour
Shrinkage-corrected log2 fold change
6
3
0
-3
-6
1
2
3
4
5
log10(mean normalised count + 1)
Up in Chili
Up in Mahmoudi
Not selected

c

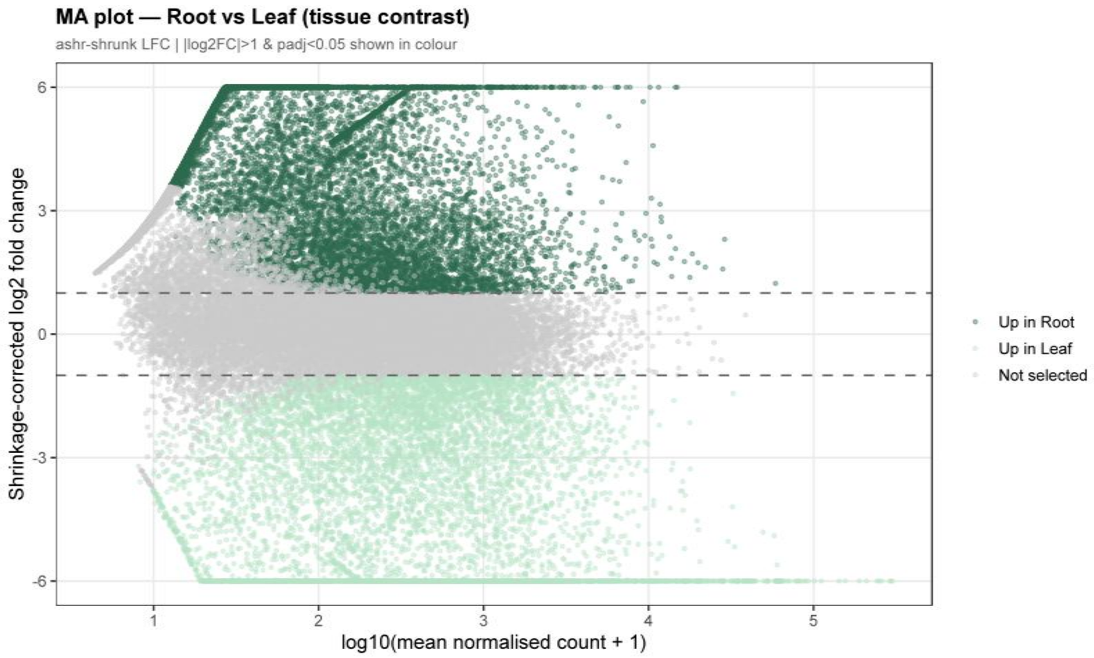
MA plot — Root vs Leaf (tissue contrast)
ashr-shrunk LFC | |log2FC|>1 & padj<0.05 shown in colour
Shrinkage-corrected log2 fold change
6
3
0
-3
-6
1
2
3
4
5
log10(mean normalised count + 1)
Up in Root
Up in Leaf
Not selected

d

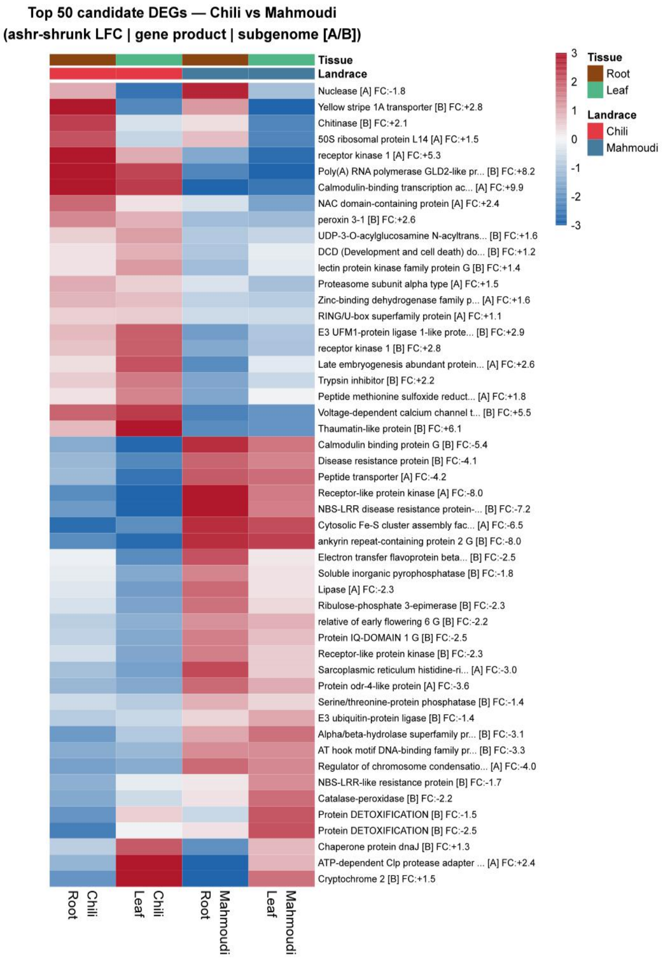

Top 50 candidate DEGs — Chili vs Mahmoudi
(ashr-shrunk LFC | gene product | subgenome [A/B])
Tissue
Landrace
Nuclease [A] FC:-1.8
Yellow stripe 1A transporter [B] FC:+2.8
Chitinase [B] FC:+2.1
50S ribosomal protein L14 [A] FC:+1.5
receptor kinase 1 [A] FC:+5.3
Poly(A) RNA polymerase GLD2-like pr... [B] FC:+8.2
Calmodulin-binding transcription ac... [A] FC:+9.9
NAC domain-containing protein [A] FC:+2.4
peroxin 3-1 [B] FC:+2.6
UDP-3-O-acylglucosamine N-acyltrans... [B] FC:+1.6
DCD (Development and cell death) do... [B] FC:+1.2
lectin protein kinase family protein G [B] FC:+1.4
Proteasome subunit alpha type [A] FC:+1.5
Zinc-binding dehydrogenase family p... [A] FC:+1.6
RING/U-box superfamily protein [A] FC:+1.1
E3 UFM1-protein ligase 1-like prote... [B] FC:+2.9
receptor kinase 1 [B] FC:+2.8
Late embryogenesis abundant protein... [A] FC:+2.6
Trypsin inhibitor [B] FC:+2.2
Peptide methionine sulfoxide reduct... [A] FC:+1.8
Voltage-dependent calcium channel t... [B] FC:+5.5
Thaumatin-like protein [B] FC:+6.1
Calmodulin binding protein G [B] FC:-5.4
Disease resistance protein [B] FC:-4.1
Peptide transporter [A] FC:-4.2
Receptor-like protein kinase [A] FC:-8.0
NBS-LRR disease resistance protein-... [B] FC:-7.2
Cytosolic Fe-S cluster assembly fac... [A] FC:-6.5
ankyrin repeat-containing protein 2 G [B] FC:-8.0
Electron transfer flavoprotein beta... [B] FC:-2.5
Soluble inorganic pyrophosphatase [B] FC:-1.8
Lipase [A] FC:-2.3
Ribulose-phosphate 3-epimerase [B] FC:-2.3
relative of early flowering 6 G [B] FC:-2.2
Protein IQ-DOMAIN 1 G [B] FC:-2.5
Receptor-like protein kinase [B] FC:-2.3
Sarcoplasmic reticulum histidine-ri... [A] FC:-3.0
Protein odr-4-like protein [A] FC:-3.6
Serine/threonine-protein phosphatase [B] FC:-1.4
E3 ubiquitin-protein ligase [B] FC:-1.4
Alpha/beta-hydrolase superfamily pr... [B] FC:-3.1
AT hook motif DNA-binding family pr... [B] FC:-3.3
Regulator of chromosome condensatio... [A] FC:-4.0
NBS-LRR-like resistance protein [B] FC:-1.7
Catalase-peroxidase [B] FC:-2.2
Protein DETOXIFICATION [B] FC:-1.5
Protein DETOXIFICATION [B] FC:-2.5
Chaperone protein dnaJ [B] FC:+1.3
ATP-dependent Clp protease adapter ... [A] FC:+2.4
Cryptochrome 2 [B] FC:+1.5
Chili Root
Chili Leaf
Mahmoudi Root
Mahmoudi Leaf
3
2
1
0
-1
-2
-3
Tissue
Root
Leaf
Landrace
Chili
Mahmoudi


e

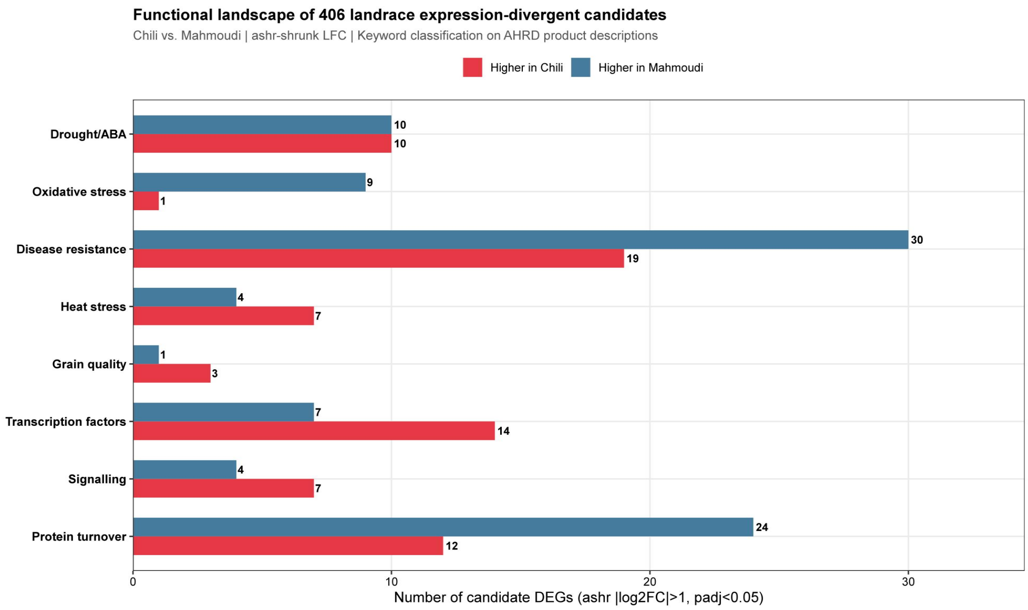


f

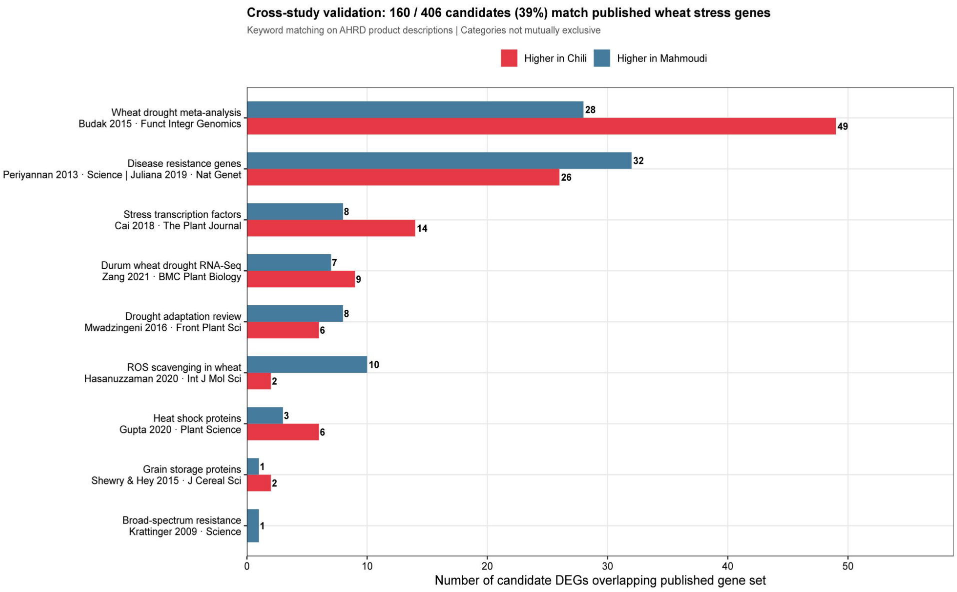


**Figure 5:** Constitutive transcriptome divergence between Tunisian durum wheat landraces. (a) PCA of variance-stabilising transformation (VST) counts for the top 500 genes by variance. PC1 (99.5% variance) separates root from leaf; PC2 (0.3% variance) resolves landrace identity. Four-way sample separation demonstrates biological coherence and absence of technical confounding. (b) MA plot of the landrace contrast: 204 Chili-elevated (red) and 202 Mahmoudi-elevated (blue) observations selected by |ashr-shrunk $\log_2$ fold change| > 1 with adjusted $P < 0.05$ as a noise filter. Note: n = 1 per landrace × tissue; this represents exploratory observation ranking, not formal hypothesis testing. (c) Tissue MA plot (positive control): 9,874 root-elevated and 7,575 leaf-elevated genes with adjusted $P < 0.05$ and |LFC| > 1, confirming DESeq2 performance. (d) Heatmap of the top 50 observations by absolute $\log_2$ fold change using VST-normalised data; colour scale shows relative expression across all four samples. (e) Functional category distribution of 406 observations across 16 manually curated AHRD-based categories (12 tested for enrichment). The six largest categories are displayed in Table 1. Disease resistance (n = 91), drought and ABA signalling (n = 38), and ROS scavenging (n = 32) are the largest classes. (f) Cross-study validation: 160 of 406 observations (39.4%) match gene families reported in nine published wheat stress studies; 246 (60.6%) are potentially novel Tunisian-landrace-specific regulatory loci.
