## Supplementary material for "Unlocking open-access genomic and transcriptomic data: the first bioinformatic exploitation of tunisian durum wheat landraces Chili and Mahmoudi": S2_statistics.txt

Figure S2 — Empirical null distribution of maximum FST
=======================================================
N permutations: 1000
Observed maximum FST: 0.833
P95 (95th percentile): 0.500
P99 (99th percentile): 0.833
Empirical p-value [(k + 1) / (N + 1)]: 0.013
  k = number of permutations with max FST >= observed (k = 12)
Interpretation: Observed max FST (0.833) equals the 99th percentile
  of 1,000 permuted values; empirical p = 0.013 under the
  (k + 1) / (N + 1) formula reported in Materials and Methods.
