## Supplementary material for "Unlocking open-access genomic and transcriptomic data: the first bioinformatic exploitation of tunisian durum wheat landraces Chili and Mahmoudi": Supplementary_Note_S1.txt

Supplementary Note S1 — Distance threshold sensitivity for FST x expression co-localisation
============================================================================================

Physical co-localisation between FST outlier windows and expression-divergent
observations was assessed at three flank distances around each outlier window:
±50 kb, ±500 kb, and ±1 Mb. The 46 permutation-significant FST outlier windows
(10-kb windows; FST >= 0.50; empirical P < 0.05; 1,000 label-shuffle permutations)
were extended by each flank and intersected against the 63,863 Svevo v1.0 GFF3
gene features; overlapping gene IDs were cross-referenced against the 406
expression-divergent observations.

Results:
  ±50 kb  : 2 of 406 observations co-localised (0.5%)
  ±500 kb : 6 of 406 observations co-localised (1.5%); 4 additional genes
            captured beyond the ±50 kb set
  ±1 Mb   : 6 of 406 observations co-localised (1.5%); count plateaus relative
            to ±500 kb, confirming all physically linked genes are recovered
            within 500 kb

On chromosome 6B, zero observations co-localised with the 6B FST hotspot at any
of the three thresholds, even though 53 expression-divergent observations map to
chromosome 6B. The interpretation that >=98.5% of expression divergence is
trans-regulatory is therefore robust to the choice of distance threshold within
the ±50 kb to ±1 Mb range.

Scripts:
  scripts/03_integration/colocalisation_50kb.py
  scripts/03_integration/colocalisation_500kb.py
