## Supplementary material for "Unlocking open-access genomic and transcriptomic data: the first bioinformatic exploitation of tunisian durum wheat landraces Chili and Mahmoudi": Supplementary_Note_S2.txt

Supplementary Note S2 — Linkage disequilibrium analysis at the chromosome 6B FST hotspot
==========================================================================================

To test whether the 53 expression-divergent observations mapping to chromosome 6B
are statistically tagged by the 6B FST hotspot, we computed pairwise r^2 between
the lead SNP at the 6B hotspot (~154.4 Mb on LT934122.1) and all SNPs within
±5 Mb on the merged biallelic VCF used for population-genomic analyses.

Methodology:
  1. The lead SNP was defined as the variant with the highest weighted FST
     within the 10-kb hotspot window on chromosome 6B.
  2. Genotypes for the lead SNP and all candidate SNPs (±5 Mb flank) were
     extracted from durum_merged_diploid.vcf.gz (n = 10 accessions; tetraploid
     genotypes converted to diploid as described in Materials and Methods).
  3. Pairwise r^2 was computed using PLINK2 --r2-unphased with --ld-window-r2 0.
  4. SNPs were binned by physical distance and median r^2 was reported per bin.

Results:
  - 50 SNPs were available within ±5 Mb of the lead SNP (sparse coverage
    reflects the small WGS panel of 10 accessions).
  - Zero of the 53 chromosome-6B expression-divergent observations had
    r^2 > 0.2 with the lead SNP.
  - Median r^2 across distance bins was <= 0.25, consistent with rapid LD decay.

Interpretation:
  The 53 chromosome-6B observations are genetically independent of the 6B
  selection signal. Physical separation (>1 Mb from the hotspot) remains the
  primary evidence for trans-regulatory divergence; the LD analysis is
  supportive but limited by sparse SNP density given n = 10 accessions.

See Figure S4 (LD decay plot) for the visual summary.
