## Supplementary figures and images for "Unlocking open-access genomic and transcriptomic data: the first bioinformatic exploitation of tunisian durum wheat landraces Chili and Mahmoudi"

### Figure_S2_Permuted_FST_Distribution.pdf

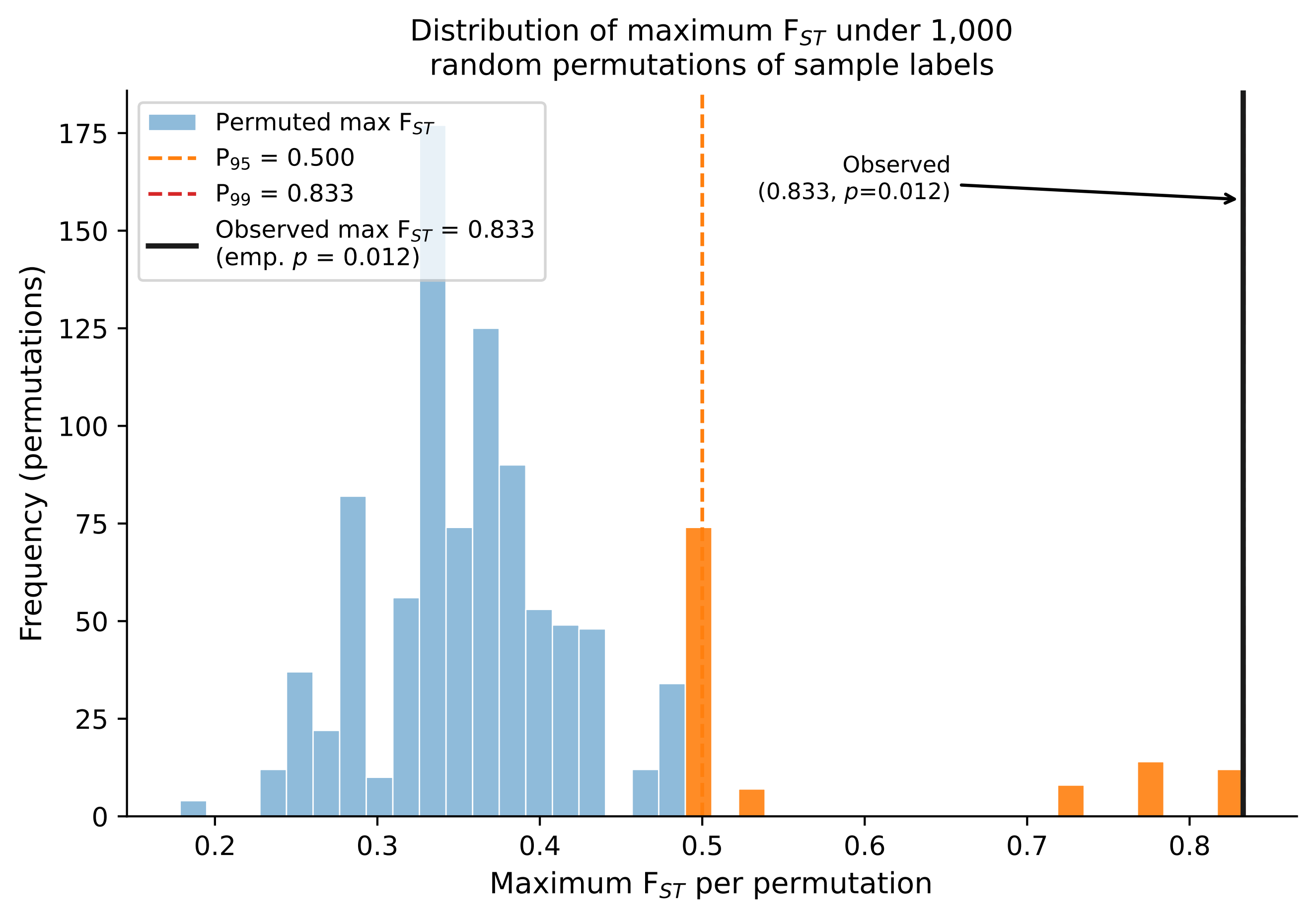

### Figure_S3_Enrichment_OR_Plot.pdf

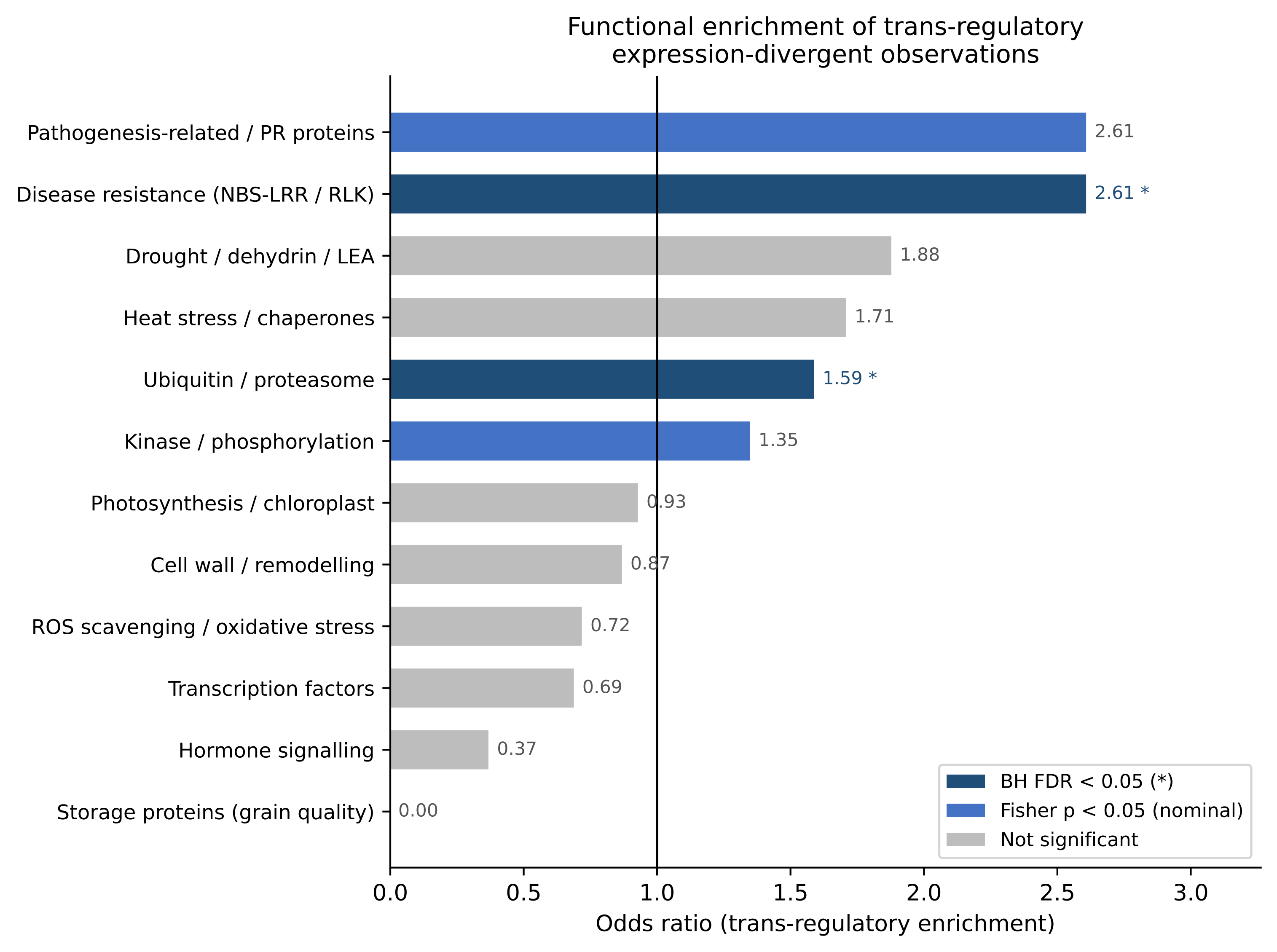

### Figure_S4_LD_Decay_6B.pdf

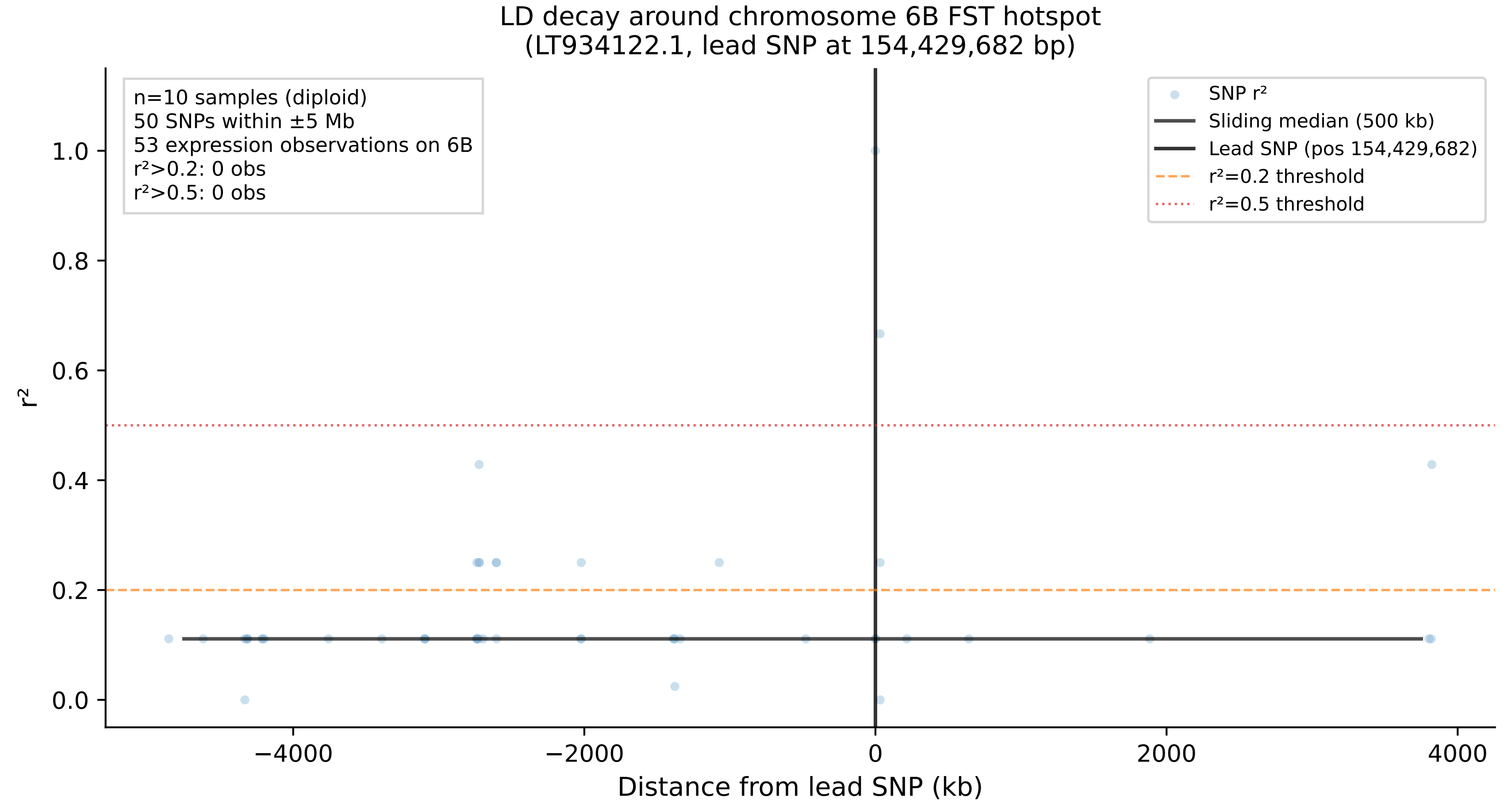

### Figure_S4_LD_Decay_6B.png

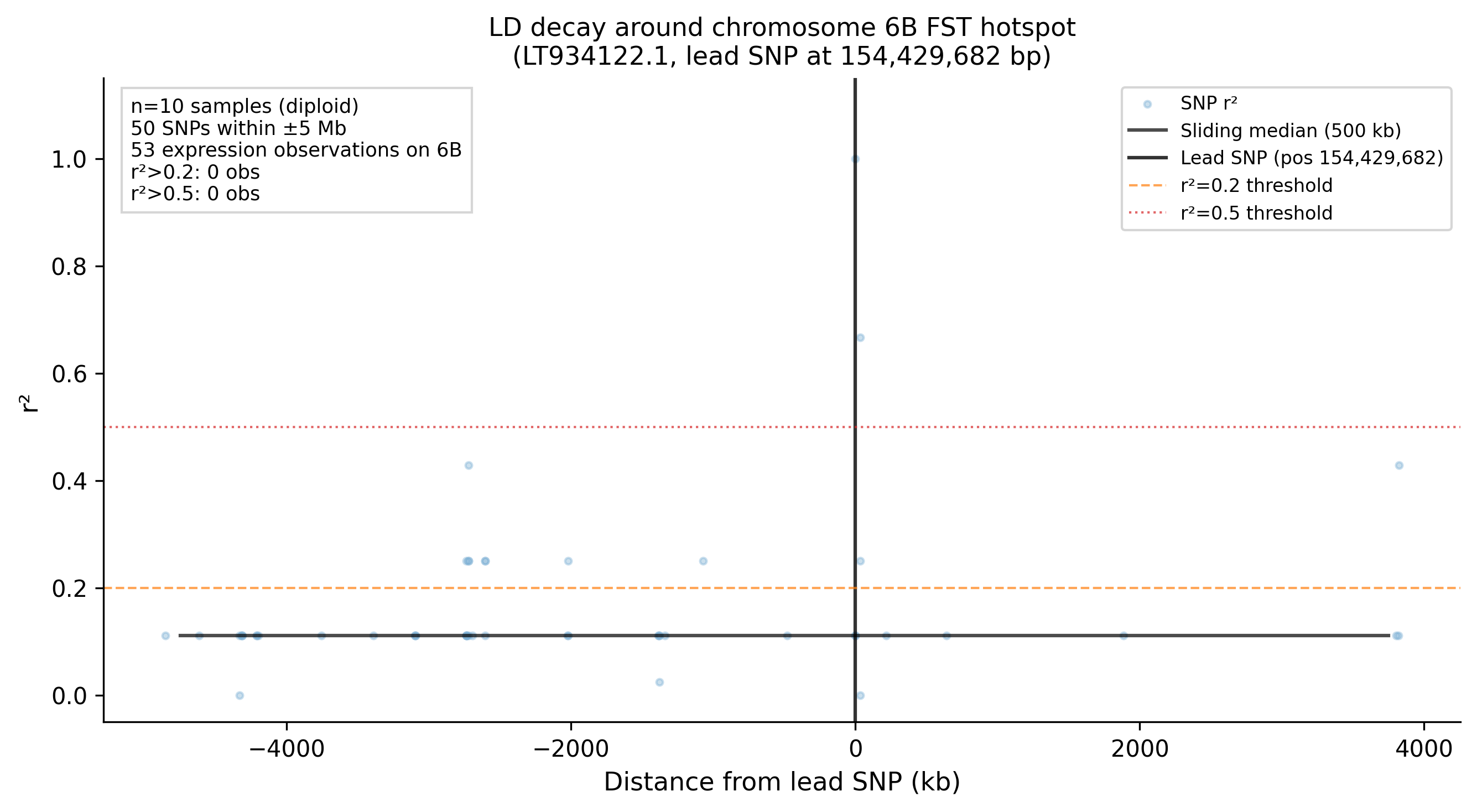
